# Small RNAs positively and negatively control transcription elongation through modulation of Rho utilization site accessibility

**DOI:** 10.1101/2024.02.02.578684

**Authors:** Kristen R. Farley, Andrew K. Buechler, Colleen M. Bianco, Carin K. Vanderpool

## Abstract

Bacteria use a multi-layered regulatory strategy to precisely and rapidly tune gene expression in response to environmental cues. Small RNAs (sRNAs) form an important layer of gene expression control and most act post-transcriptionally to control translation and stability of mRNAs. We have shown that at least five different sRNAs in *Escherichia coli* regulate the cyclopropane fatty acid synthase (*cfa*) mRNA. These sRNAs bind at different sites in the long 5’ untranslated region (UTR) of *cfa* mRNA and previous work suggested that they modulate RNase E-dependent mRNA turnover. Recently, the *cfa* 5’ UTR was identified as a site of Rho-dependent transcription termination, leading us to hypothesize that the sRNAs might also regulate *cfa* transcription elongation. In this study we find that a pyrimidine-rich region flanked by sRNA binding sites in the *cfa* 5’ UTR is required for premature Rho-dependent termination. We discovered that both the activating sRNA RydC and repressing sRNA CpxQ regulate *cfa* primarily by modulating Rho-dependent termination of *cfa* transcription, with only a minor effect on RNase E-mediated turnover of *cfa* mRNA. A stem-loop structure in the *cfa* 5’ UTR sequesters the pyrimidine-rich region required for Rho-dependent termination. CpxQ binding to the 5’ portion of the stem increases Rho-dependent termination whereas RydC binding downstream of the stem decreases termination. These results reveal the versatile mechanisms sRNAs use to regulate target gene expression at transcriptional and post-transcriptional levels and demonstrate that regulation by sRNAs in long UTRs can involve modulation of transcription elongation.

**Importance:** Bacteria respond to stress by rapidly regulating gene expression. Regulation can occur through control of messenger RNA (mRNA) production (transcription elongation), stability of mRNAs, or translation of mRNAs. Bacteria can use small RNAs (sRNAs) to regulate gene expression at each of these steps, but we often do not understand how this works at a molecular level. In this study, we find that sRNAs in *Escherichia coli* regulate gene expression at the level of transcription elongation by promoting or inhibiting transcription termination by a protein called Rho. These results help us understand new molecular mechanisms of gene expression regulation in bacteria.

## Introduction

Small regulatory RNAs in bacteria, archaea, and eukaryotes promote rapid changes in gene expression and subsequent changes in cell physiology or behavior (1). Many small RNAs (sRNAs) exert their regulatory effects by base pairing with target mRNAs (2) via short, semi-complementary interactions (3). In *Escherichia coli* and related bacteria, sRNAs require the RNA chaperone Hfq for stability and mRNA target regulation (4). Many Hfq-dependent sRNAs base pair near the ribosome binding site (RBS) on target mRNAs to modulate translation and (indirectly) mRNA stability. Some sRNAs directly modulate mRNA stability by blocking ribonuclease E (RNase E) cleavage sites (4, 5). Although bacterial sRNAs are best known as post-transcriptional regulators, recent work has revealed that some sRNAs modulate transcription elongation through interactions in long 5’ untranslated regions (UTRs) of mRNA targets (6, 7). About 25% of *E. coli* mRNAs possess 5’ UTRs longer than 80 nt and the levels of approximately half of these long-UTR mRNAs are altered by bicyclomycin, an inhibitor of transcription termination factor Rho (7). These observations suggest that Rho-dependent termination events in long 5’ UTRs may be common and that there may be more sRNAs that regulate target mRNAs by modulating transcription elongation.

Rho is a hexameric protein that functions as an essential transcription termination factor in most bacteria (8). Rho binds to nascent RNAs at sequences called Rho utilization (*rut*) sites and hydrolyzes ATP to translocate toward RNA polymerase (RNAP) and the transcription elongation complex. Rho destabilizes the elongation complex, leading to transcription termination (8). Active translation shields mRNAs from Rho-dependent transcription termination within coding sequences but uncoupling of transcription and translation can leave nascent mRNAs susceptible to premature Rho-dependent termination (8). While it is evident how sRNA-mediated translational repression renders target mRNAs susceptible to both RNases and Rho, it has been unclear how sRNA regulators that do not directly affect translation modulate transcription elongation or mRNA stability (6). Nevertheless, there are several examples of such translation-independent regulation. The sRNAs DsrA, ArcZ, and RprA prevent Rho-dependent termination within the *rpoS* mRNA 5’ UTR (7) by a mechanism proposed to involve disruption of Rho binding or translocation. The sRNA SraL represses termination within the *rho* mRNA, which is autoregulated (9, 10). While the mechanism is unclear, SraL is thought to induce *rho* mRNA structural changes to make a *rut* site inaccessible (10).

The *cfa* mRNA in *E. coli* is regulated positively and negatively by sRNAs that base pair at different sites within a 212-nt 5’ UTR (11, 12). The *cfa* gene encodes cyclopropane fatty acid (CFA) synthase that installs cyclopropane rings in unsaturated membrane phospholipids (13). Under acidic and oxidative stress conditions, CFAs are thought to promote cell survival by stabilizing the cell membrane and reducing membrane permeability (13, 14). The sRNA RydC is a robust activator of *cfa* (12), while CpxQ (15) represses *cfa* (11). Previous studies suggested that these sRNAs regulate *cfa* by modulating RNase E-mediated *cfa* mRNA decay (11, 12, 16). However, a recent study found evidence for Rho-dependent transcription termination in the 5’ region of the *cfa* coding sequence (CDS) (17), prompting us to investigate whether sRNAs regulate *cfa* through modulating Rho-dependent termination.

In this study we found that Rho limits transcription elongation from a σ^70^ promoter that produces a long isoform of *cfa* mRNA. The sRNA binding sites within the *cfa* 5’ UTR flank a pyrimidine-rich region that has hallmarks of a *rut* site, and this region is critical for Rho-dependent *cfa* regulation. The activating sRNAs ArrS and RydC and the repressing sRNAs CpxQ and GcvB all require Rho to carry out regulation of *cfa*. Our data are consistent with a model where sRNAs primarily act via modulation of Rho-dependent termination and not by control of RNase E-dependent turnover of *cfa* mRNA. This work reveals that sRNAs are not solely post-transcriptional regulators, but instead that some regulate their mRNA targets in a translation-independent manner at the level of transcription elongation.

## Results

### Rho regulates transcription of the long *cfa* mRNA isoform

The *cfa* σ^70^ promoter produces a long mRNA isoform with a 212-nt 5’ UTR, and a σ^S^ promoter produces a short isoform with a 34-nt 5’ UTR (18) (Figure 1A). To determine if either isoform is subject to premature Rho-dependent termination, we measured the activity of *cfa* reporter fusions in *rho*^+^ and *rho*-R66S mutant strains. (Rho-R66S has a transcription termination defect due to reduced RNA binding (19).) Rho autoregulates its own transcription, and activity of a *rho*’-’*lacZ* fusion was higher in the *rho*-R66S mutant compared to the *rho^+^*strain (Figure S1), indicating that the *rho-*R66S mutation has the expected effect on Rho activity (20). The activity of a *cfa*-Short (34-nt 5’ UTR) translational fusion was similar in both *rho^+^*and *rho*-R66S backgrounds, whereas activity of the *cfa*-Long (212-nt 5’ UTR) fusion was higher in the *rho*- R66S mutant (Figure 1B).

**Figure 1.**
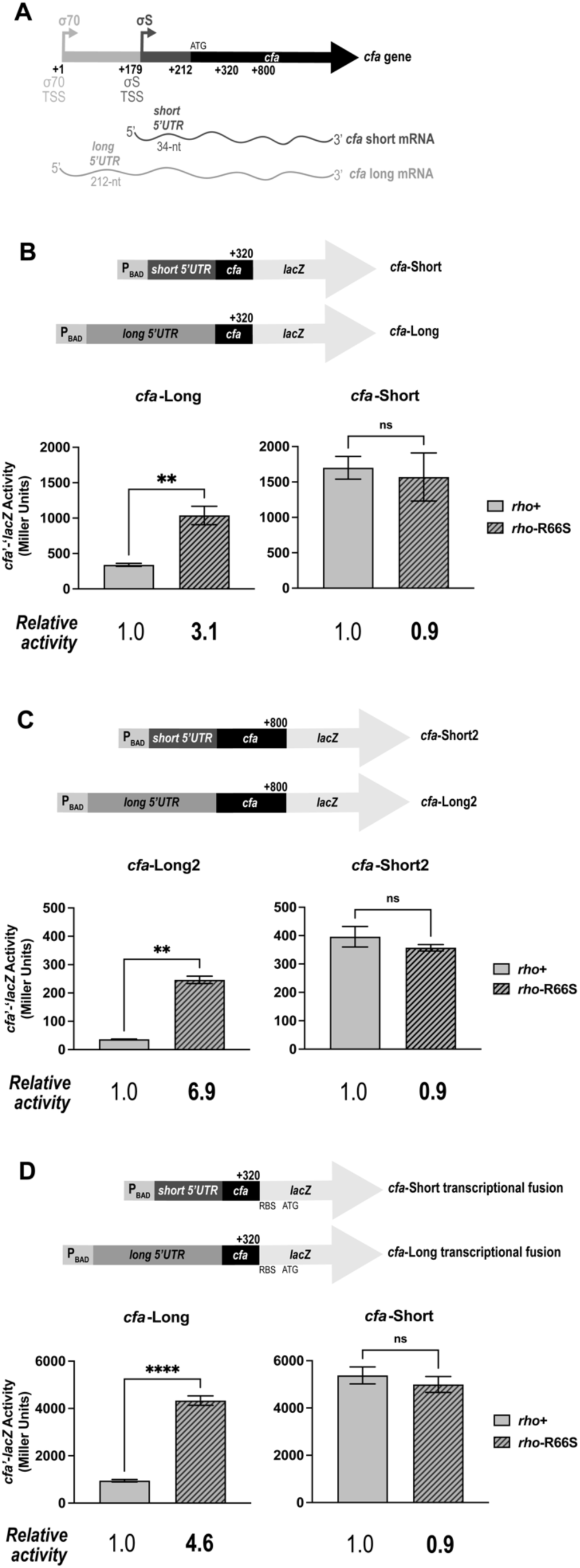
The long 5’ UTR of *cfa* mRNA is required for Rho-dependent regulation. **A.** The long *cfa* mRNA isoform contains a 212- nt 5’ UTR (*cfa*-Long), while the short *cfa* mRNA isoform contains a 34-nucleotide 5’ UTR (*cfa*-Short). **B.** *cfa*-Long and *cfa*-Short translational fusions contain the UTRs and the first 36 codons of the *cfa* coding sequence (to position +320-nt) fused to the 9^th^ codon of the *lacZ* CDS (top). Both fusions are controlled by an arabinose-inducible promoter (P_BAD_). β-galactosidase activity of fusion strains was measured at mid-exponential phase. Error bars represent the standard deviations of three biological replicates and statistical significance was determined using two-tailed Welch’s t-tests by comparing the fusion activity in the *rho*^+^ strain versus the *rho*-R66S mutant (**P < 0.05). **C.** *cfa*-Long2 and *cfa*-Short2 fusions contain the UTR regions with the first 196 codons (to +800-nt) of the *cfa* CDS fused to the 9^th^ codon of the *lacZ* CDS (top). β- galactosidase activity was measured and data analyzed as described in B. **D.** *cfa*- Long and *cfa*-Short transcriptional fusion constructs contain the same fragments as the corresponding translational fusions in B. β-galactosidase activity was measured and data analyzed as described in B. (****P < 0.0001).

In a recent transcriptome study, multiple *cfa* mRNA 3’ ends were identified (17). One was at position +59 (relative to the long isoform start site) and another was at position +330 within the *cfa* CDS. A third 3’ end at +446 was bicyclomycin-dependent, suggesting a putative Rho-dependent termination event at or near this site (17). We hypothesized that Rho binds to a *rut* site in the *cfa* long 5’ UTR and terminates transcription within the 5’ UTR or CDS. To determine if sequences in the *cfa* CDS influence regulation, we constructed *cfa’-’lacZ* fusions that extend to position +800 (*cfa*-Long2, *cfa*-Short2, Figure 1C). Activity of the *cfa*-Short2 fusion was similar in *rho^+^* and *rho*-R66S mutant strains, while activity of the *cfa*-Long2 fusion was significantly higher in the *rho*-R66S mutant (Figure 2C). The fold-change in activity in *rho^+^* compared to *rho-*R66S is higher for the *cfa-*Long2 than for the *cfa*-Long fusion (compare Figures 1B and 1C). Activity of analogous transcriptional fusions showed the same result. There was no Rho-dependent regulation of a *cfa*-Short transcriptional fusion and strong regulation of a *cfa*-Long transcriptional fusion (Figure 1D).

**Figure 2.**
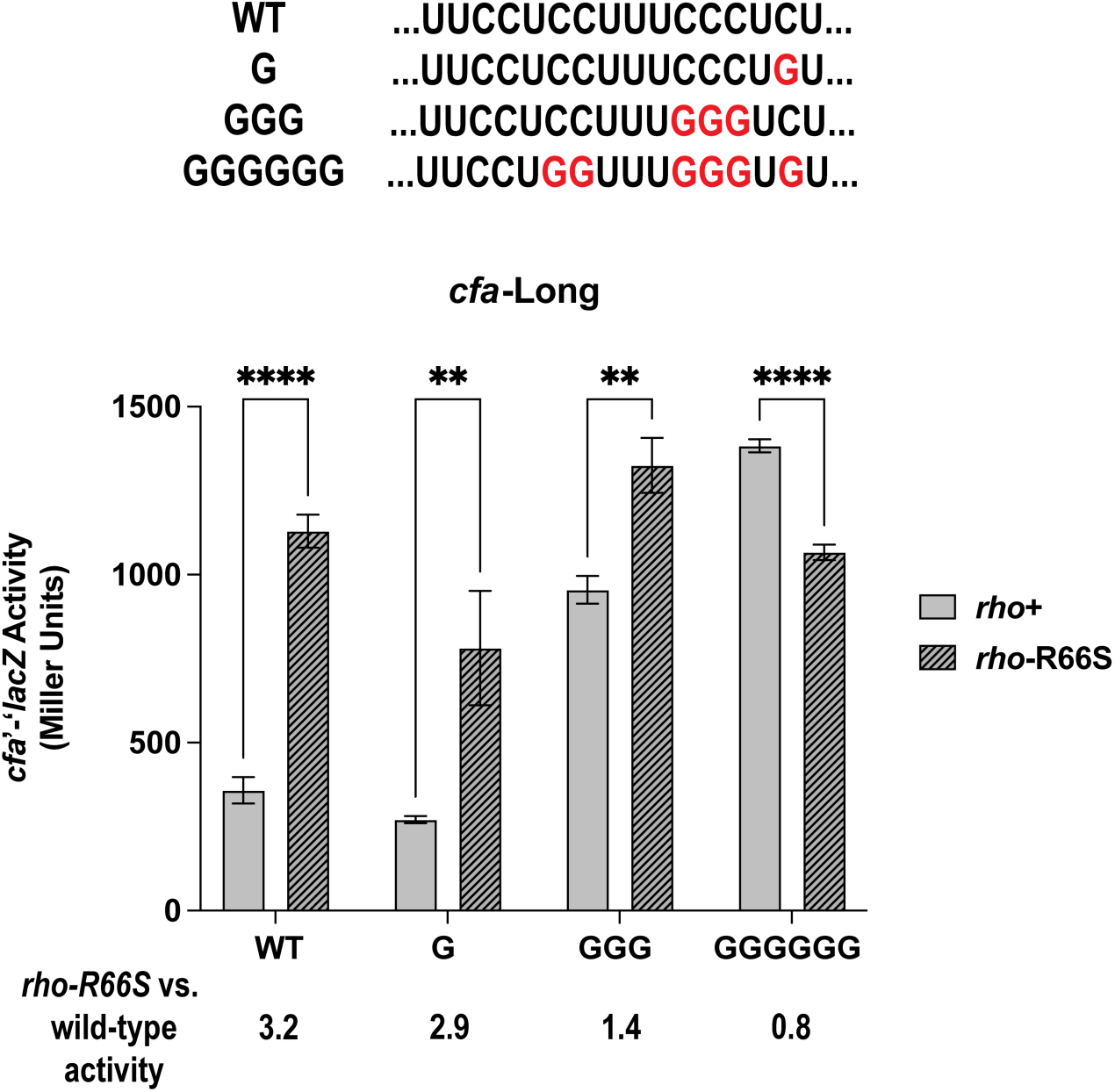
A pyrimidine-rich tract in the *cfa* long 5’ UTR is required for Rho-dependent regulation. The CU-rich tract identified within the *cfa* long 5’ UTR is shown. Variant fusions have C:G substitutions as shown in red font. The β-galactosidase activity of each fusion was tested in *rho^+^* and *rho-*R66S mutant strain backgrounds. β-galactosidase activity is expressed in Miller Units and the activity was measured at mid-exponential phase. Error bars represent the standard deviations of three biological replicates and statistical significance was determined using multiple unpaired Welch’s t-tests by comparing the activity in the wild-type strain versus the *rho*-R66S mutant (****P < 0.0001).

### A pyrimidine-rich tract in the *cfa* 5’ UTR is required for Rho-dependent regulation

While *rut* sites have no consensus sequence, they typically comprise ∼80-nt C-rich/G-poor unstructured regions (6). Some *rut* sites have pyrimidine-rich tracts that promote Rho-dependent transcription termination (21), and we identified a 16-nt CU-rich region between +86 and +101 of the *cfa* long 5’ UTR. To test the role of this region in Rho-dependent regulation of *cfa*, we measured activity of *cfa*-Long fusion variants with C to G substitutions. While a single substitution (G, Figure 2) had a minimal impact on Rho-dependent regulation, three substitutions (GGG, Figure 2) resulted in strongly increased activity in the *rho^+^*strain, suggesting the GGG variant is less susceptible to Rho-dependent termination. Six substitutions (GGGGGG, Figure 2), resulted in a further increase in fusion activity in the *rho*^+^ strain, and activity of this fusion was not increased in the *rho*-R66S compared to the *rho^+^* background. This result suggests that Rho cannot promote premature termination of the GGGGGG variant. These data are consistent with the hypothesis that the 16-nt CU-rich region is part of a larger *rut* site required for Rho-dependent regulation of *cfa*.

Alignment of *cfa* 5’ UTR sequences from 10 different bacterial species belonging to the Enterobacteriaceae (Figure S2A) revealed strong conservation of the CU-rich region (Figure S2A). Since *cfa* in *S. enterica* is regulated by some of the same factors as *E. coli cfa*, including RNase E and the sRNA RydC (12, 16), we tested whether *S. enterica cfa* is regulated by Rho. The activity of *S. enterica cfa*-Long and *cfa*-Short translational fusions (*Stm* fusions, Figure S2B) in the *E. coli* wild-type and *rho*-R66S mutant backgrounds showed that *Stm cfa*-Long fusion activity is higher in the *rho*-R66S mutant compared to the wild-type strain (Figure S2B). These data suggest that Rho-dependent regulation of *cfa* is conserved between *E. coli* and *Salmonella*.

To further characterize the region required for Rho-dependent termination, we measured activity of truncated *cfa*-Long transcriptional fusions (Figure 3). While basal activities of the fusions varied, fusions containing the 5’ UTR with sequences at or beyond +138 displayed more than 2-fold higher levels of activity in the *rho*-R66S compared to the *rho^+^*strain (Figure 3). Shorter fusions (+58, +78, +98, +118) showed little or no Rho-dependence (Figure 3). The CU-rich region is located between positions +85 and +102 (Figure 4A). These data indicate that sequences downstream of this region are also required for efficient Rho-dependent termination in the *cfa* 5’ UTR. Thus, these data suggest that the CU-rich region and sequences immediately downstream constitute a *rut* site required for Rho-dependent termination of *cfa* transcription.

**Figure 3.**
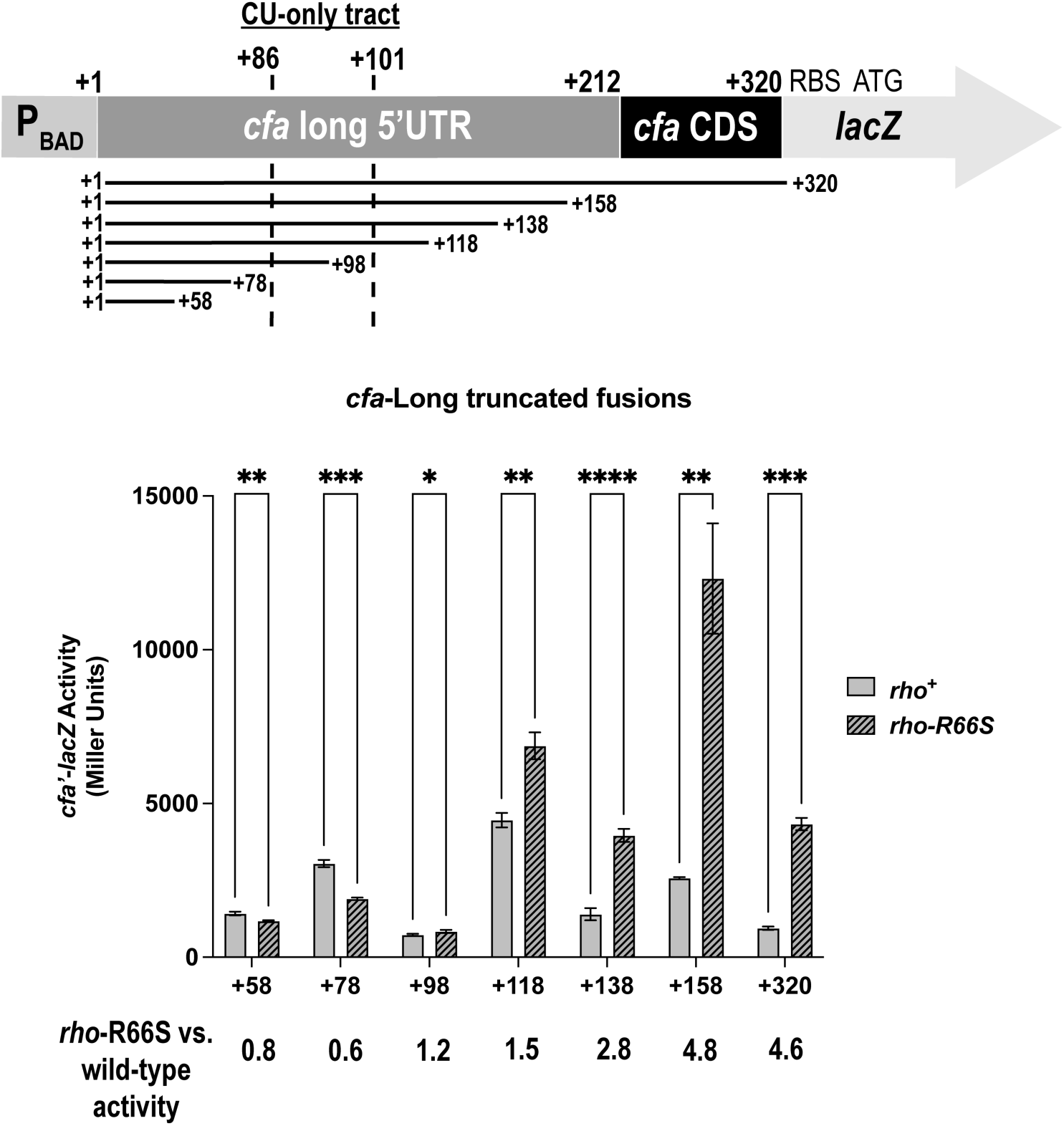
The *cfa* long 5’ UTR contains all sequences needed for Rho-dependent regulation. Six *cfa*-Long transcriptional fusions that were truncated from the 3’ end were constructed (top). The β-galactosidase activity of each truncated fusion was tested in the *rho^+^* and *rho-*R66S mutant strain background (bottom). β-galactosidase activity is expressed in Miller Units and the activity was measured at mid-exponential phase. Error bars represent the standard deviations of three biological replicates and statistical significance was determined using multiple unpaired Welch’s t-tests by comparing the activity in the wild-type strain versus the *rho*-R66S mutant (****P < 0.0001).

**Figure 4.**
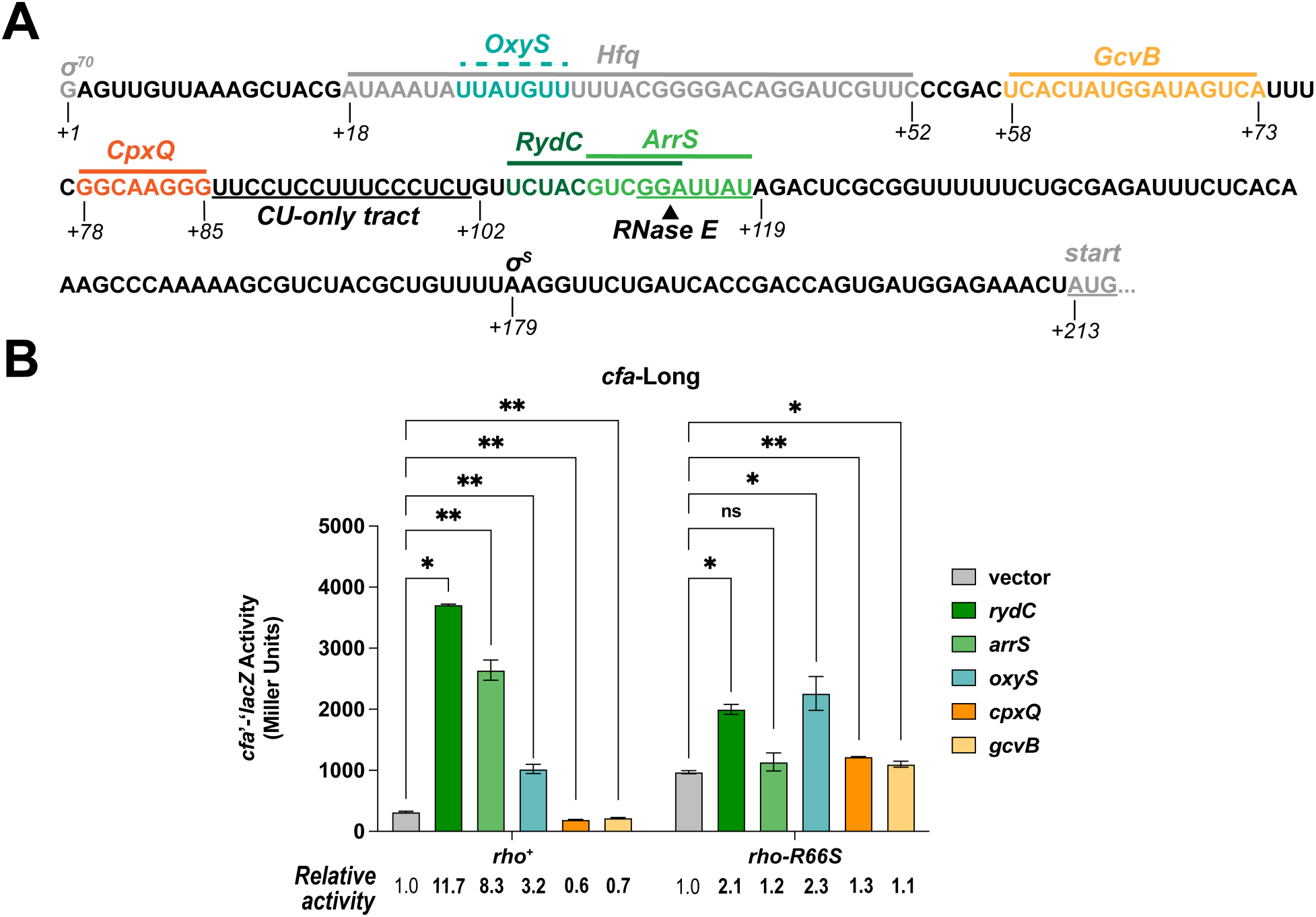
Activating and repressing sRNAs modulate Rho-dependent regulation of *cfa* mRNA. **A.** The *cfa* σ^70^ transcription start site position is +1 and binding sites are indicated with solid lines (for validated binding sites) or dotted lines (for predicted sites) above the sequence. The predicted RNase E cleavage site and CU-rich region are underlined. The arrow indicates the predicted RNase E cleavage point. **B.** β-galactosidase activity of the *cfa*-Long translational fusion in the *rho*^+^ or *rho-* R66S mutant strain background in the presence of vector control or plasmids producing the sRNAs RydC, ArrS, OxyS, CpxQ, or GcvB. β-galactosidase activity is expressed in Miller Units and the activity was measured at mid-exponential phase after one hour of sRNA induction. Error bars represent the standard deviations of three biological replicates and statistical significance was determined using a multiple unpaired Welch’s t-tests (**P < 0.05).

### Activating and repressing sRNAs modulate Rho-dependent *cfa* regulation

The long *cfa* mRNA isoform is regulated by several Hfq-dependent sRNAs. RydC, ArrS, and OxyS activate *cfa*, while CpxQ and GcvB repress *cfa* (11). RydC and ArrS bind to overlapping sites (Figure 4A) and previous work suggested that they occlude an RNase E cleavage site and impair degradation of *cfa* mRNA (11, 12, 16). CpxQ binds upstream of the RydC binding site (Figure 4A) and we reported that CpxQ regulates *cfa* by a mechanism requiring the same RNase E cleavage site (11). RydC and ArrS bind immediately downstream of the CU-rich region (within the window required for efficient Rho-dependent termination) while CpxQ binds directly upstream of this region (Figure 4A). We hypothesized that in addition to modulating RNase E-mediated degradation, RydC/ArrS and CpxQ modulate Rho-dependent termination of *cfa* transcription.

We measured activity of the *cfa*-Long fusion in *rho*^+^ and *rho*-R66S mutant strains carrying a vector control or sRNA expression plasmid (Figure 4B). These sRNAs do not regulate the *cfa*-Short fusion (11). RydC and ArrS activate *cfa*-Long fusion activity in the *rho*^+^ strain, while CpxQ represses it. In the *rho*-R66S mutant, the magnitude of RydC-dependent activation of *cfa* is strongly diminished, and ArrS-dependent activation and CpxQ-dependent repression are completely abolished in this background (Figure 4B). Northern blot analysis showed that RydC and CpxQ levels are equivalent in *rho*^+^ and *rho*-R66S mutant strains (Figure S3). We observed the same patterns of sRNA- and Rho-dependent regulation for the *cfa*-Long2 translational fusion and *cfa*-Long transcriptional fusion (Figure S4). OxyS modestly activates the *cfa-*Long fusion (11) and the predicted OxyS binding site is far upstream of the CU-rich region (Figure 4A). OxyS-mediated activation is only slightly diminished in the *rho*-R66S mutant compared to the *rho*^+^ strain (Figure 4B) suggesting that OxyS uses a distinct mechanism to activate *cfa*. GcvB binds upstream of the CpxQ binding site (11) and modestly represses the *cfa-*Long (Figure 4A); repression is abolished in the *rho*-R66S mutant strain (Figure 4B).

### RydC and CpxQ impact *cfa* transcription elongation

To further examine the roles of RydC and CpxQ in modulating premature termination, we used RT-qPCR to measure the relative levels of different portions of *cfa* mRNA in Δ*rydC* Δ*cpxQ* mutant cells expressing either RydC or CpxQ. Probes were specific for the *cfa* 5’ UTR, the 5’ CDS, and the 3’ CDS (CDS). Figure 5A illustrates how ratios of the PCR products change with increased or decreased termination. When RydC was produced, the *cfa* 5’ UTR:CDS ratio was slightly decreased (Figure 5B, left panel), and the *cfa* 5’ CDS:CDS ratio was significantly decreased relative to the control (Figure 5B, right panel), consistent with RydC inhibition of premature termination. The data suggest that RydC promotes elongation past the 5’ region of the CDS and into the 3’ region of the *cfa* CDS. In CpxQ-producing cells, the *cfa* 5’ UTR:CDS ratio is increased, consistent with a CpxQ-mediated increase in termination (Figure 5B). There is no significant change in 5’ CDS:CDS ratios compared to the control in CpxQ-producing cells. These data suggest that CpxQ modulates termination in the region between the 5’ UTR and 5’ CDS probes. These data support the model that CpxQ and RydC modulate *cfa* transcription elongation.

**Figure 5.**
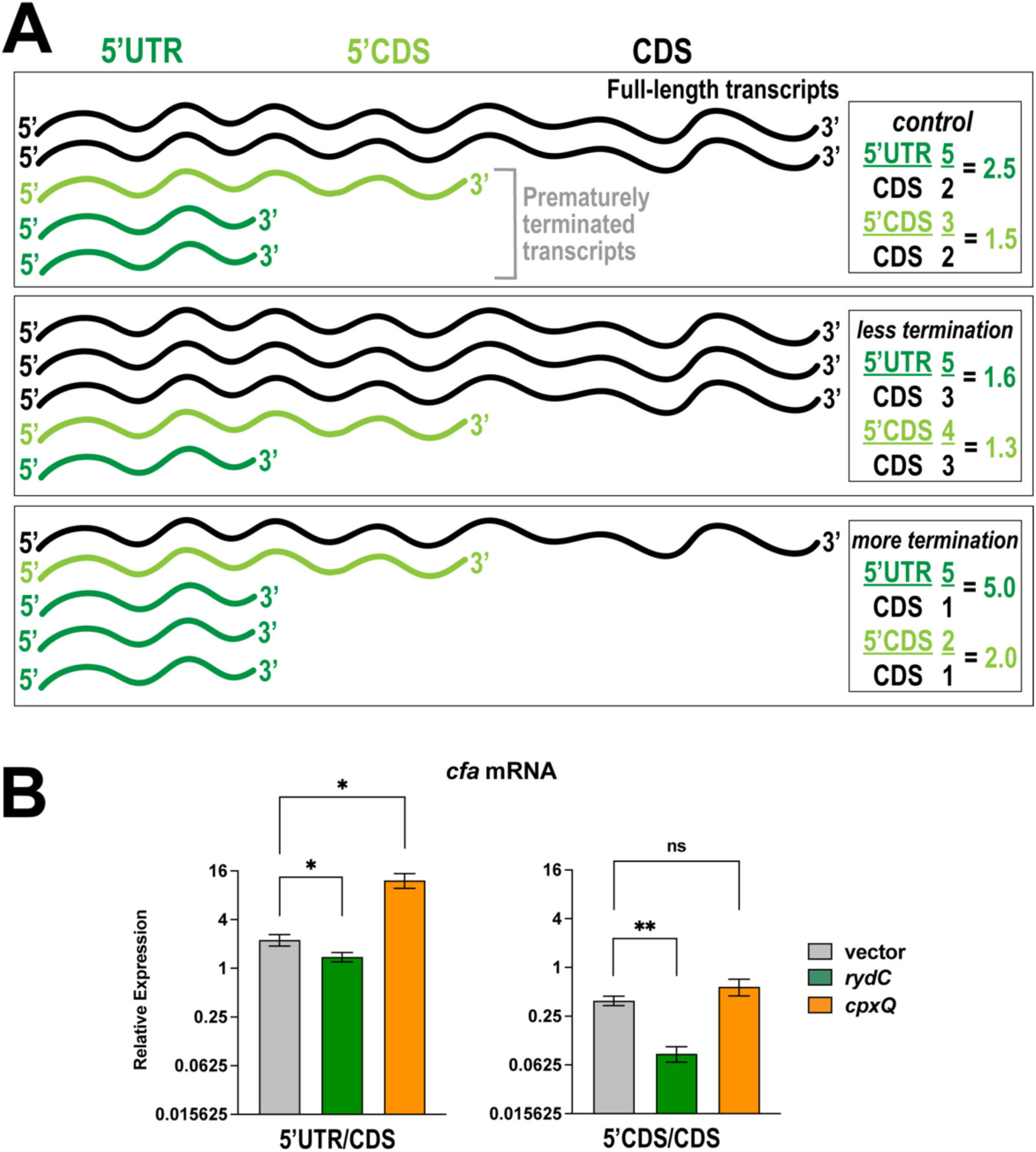
RydC and CpxQ modulate production of full-length *cfa* mRNA. **A.** Predictions for ratios of different regions of a transcript subject to Rho-dependent termination with increased or decreased efficiency of Rho-dependent termination. **B.** RT-qPCR analysis was used to measure the relative levels of different regions of the *cfa* mRNA in Δ*cpxQ* Δ*rydC* mutant cells containing an empty vector or plasmid expressing either RydC or CpxQ. The relative levels of three different regions of *cfa* mRNA were measured including the long 5’ UTR (5’ UTR), the 5’ portion of the CDS (5’ CDS), and the 3’ portion of the CDS (CDS). Error bars represent the standard deviations of three biological replicates and statistical significance was determined using multiple unpaired Welch’s t-tests (**P < 0.05).

### CpxQ binding to a stem-loop in the *cfa* 5’ UTR governs *rut* site accessibility

Other sRNAs that modulate Rho-dependent termination within 5’ UTRs are thought to inhibit Rho binding to the mRNA target, leading to increased transcription elongation (7, 10). In contrast, CpxQ increases Rho-dependent termination in the *cfa* 5’ UTR (Figure 5B). We hypothesized that CpxQ binding to *cfa* mRNA alters its secondary structure to increase the accessibility of the *rut* site. We found a putative stem-loop structure from +78 to +97 of the 5’ UTR (Figure 6A). The 5’ half of the stem contains the CpxQ binding site, while the loop and 3’ half of the stem encompass the pyrimidine-rich region that is critical for Rho-dependent termination (Figure 6A). We predicted that formation of this stem-loop occludes a critical portion of the *rut* site and limits Rho-dependent termination. Binding by CpxQ would prevent formation of the stem, leaving the *rut* site more accessible and leading to increased Rho-dependent termination.

**Figure 6.**
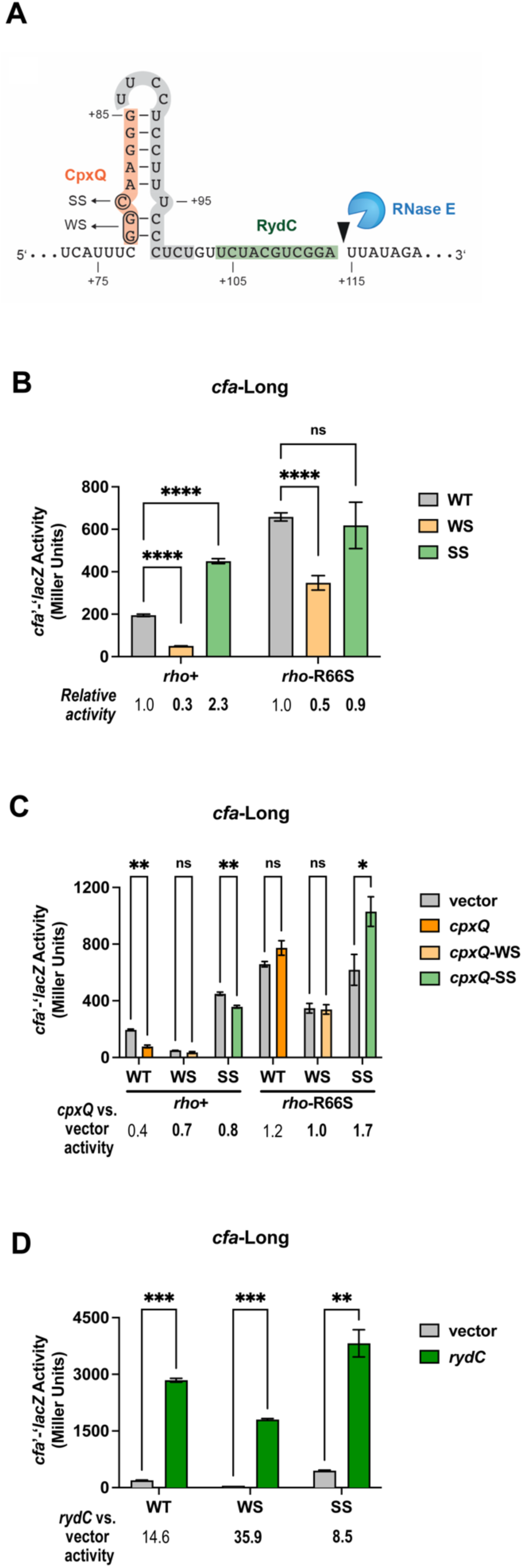
CpxQ enhances *rut* site accessibility by preventing stem-loop formation within the *cfa* long 5’ UTR. **A.** The stem-loop structure in the *cfa* long 5’ UTR that encompasses the CU- rich portion of the putative *rut* site (shaded gray) and the CpxQ binding site (shaded orange). The RydC binding site (shaded green) and a putative RNase E cleavage site are indicated. **B.** *cfa*-Long translational fusion variants: weak stem (WS) disrupts the stem-loop via two G:C substitutions and strong stem (SS) reinforces the stem-loop via a C:A substitution. Activity of wild-type (WT) *cfa*-Long, WS, and SS variant fusions was tested in *rho^+^* and *rho-*R66S mutant strain backgrounds. β-galactosidase activity is expressed in Miller Units and the activity was measured at mid-exponential phase. Error bars represent standard deviations of three biological replicates; statistical significance was determined using multiple unpaired Welch’s t-tests (****P < 0.0001). **C.** Activity of WT, WS, and SS *cfa*-Long fusions in *rho^+^* and *rho-*R66S mutant strain backgrounds carrying vector control or plasmids producing CpxQ, CpxQ-WS, or CpxQ-SS. β-galactosidase activity was measured as described in B after one hour of sRNA induction. Data were analyzed as described in B. (**P < 0.05). **D.**β- galactosidase activity of WT, WS, and SS *cfa*- Long fusions in *rho^+^* and *rho-*R66S mutant strain backgrounds carrying vector control or RydC- producing plasmids. β-galactosidase activity was measured as described in B after one hour of sRNA induction. Data were analyzed as described in B. (***P < 0.001).

To test the role of this stem in Rho-dependent termination of *cfa*, we designed two *cfa*- Long translational fusions to either weaken or strengthen the stability of the stem (Figure 6A). The *cfa*-Long-G78C-G79C fusion, referred to as *cfa*-WeakStem (*cfa-*WS), should disrupt stem formation and increase Rho-dependent termination. The *cfa*-Long-C80A fusion referred to as *cfa*-StrongStem (*cfa*-SS), should strengthen the stem, leading to decreased Rho-dependent termination. We measured activity of *cfa*-Long, *cfa*-WS, and *cfa*-SS, and found that *cfa*-WS exhibited a significant decrease in activity compared with *cfa*-Long (Figure 6B), consistent with the model that the mutations increased Rho access to the *rut* site and increased termination. In contrast, the *cfa*-SS fusion displayed an ∼2-fold increase in activity compared with *cfa*-Long (Figure 6B), consistent with reduced Rho access to the *rut* site and reduced termination. Consistent with the hypothesis that differences in reporter activity are due to differences in Rho-dependent termination, the impact of the stem-weakening and stem-strengthening mutations was either attenuated or abolished in the *rho*-R66S mutant background (Figure 6B).

We hypothesized that the WS and SS mutations would blunt the impact of CpxQ on *cfa.* CpxQ variants with compensatory mutations that restore base pairing with *cfa-*WS (CpxQ-WS) and *cfa-*SS (CpxQ-SS) were expressed in *rho*+ and *rho-*R66S backgrounds. In contrast with the wild-type interaction, which inhibits *cfa* fusion activity, there is no statistically significant repression by CpxQ-WS or CpxQ-SS on the relevant fusions in the *rho*^+^ background. In the *rho-* R66S background, there was no CpxQ-mediated repression, though we did observe a small CpxQ-SS-mediated activation of *cfa*-SS that we cannot explain (Figure 6C).

The RydC binding site is downstream of the CU-rich stem-loop, within the region (+1 to +138) important for Rho-dependent regulation of *cfa* (Figure 3). We hypothesize that RydC directly occludes Rho binding by blocking part of the *rut* site that lies downstream of the CU-rich stem-loop. If this is true, we expected that RydC-dependent activation of *cfa* is independent of formation of the stem-loop. To test this hypothesis, we measured activity of *cfa-*Long, *cfa-*WS, and *cfa-*SS fusions in strains carrying a vector control or *rydC* expression plasmid. As predicted, RydC strongly activated all three fusions (Figure 6D). Together, these data are consistent with the model that repressing and activating sRNAs regulate *cfa* by modulating access of Rho to different portions of a *rut* site in the *cfa* 5’ UTR.

### Interplay between Rho and RNase E degradosome in regulation of *cfa*

Previous studies in *S. enterica* and *E. coli* showed that *cfa* mRNA turnover is regulated by RNase E cleavage within the 5’ UTR (12, 16) and suggested that RydC and CpxQ inhibit or enhance cleavage, respectively (11). We measured *cfa*-Long fusion activity in strains with the *rne131* allele that encodes a degradosome-deficient RNase E variant (22), which cannot carry out many sRNA-dependent regulatory effects (23–25). Activity of the fusion was similar in *rne^+^* and *rne131* mutant backgrounds, and was approximately 3-fold higher in the *rho-*R66S mutant. Activity in the *rho-*R66S *rne131* double mutant was similar to the *rho-*R66S mutant (Figure 7A), suggesting that degradosome-mediated decay does not contribute significantly to *cfa* regulation. RydC and CpxQ regulate the *cfa*-Long fusion similarly in wild-type and *rne131* single mutant strains (Figure 7B). In the *rho-*R66S background, CpxQ-dependent repression and RydC-dependent activation are both diminished and there is no substantial difference in sRNA-mediated regulation between *rho*-R66S mutant and *rho*-R66S *rne131* double mutant strains (Figure 7B). These results suggest that RydC and CpxQ primarily regulate *cfa* by modulating Rho-dependent termination and not degradosome activity.

**Figure 7.**
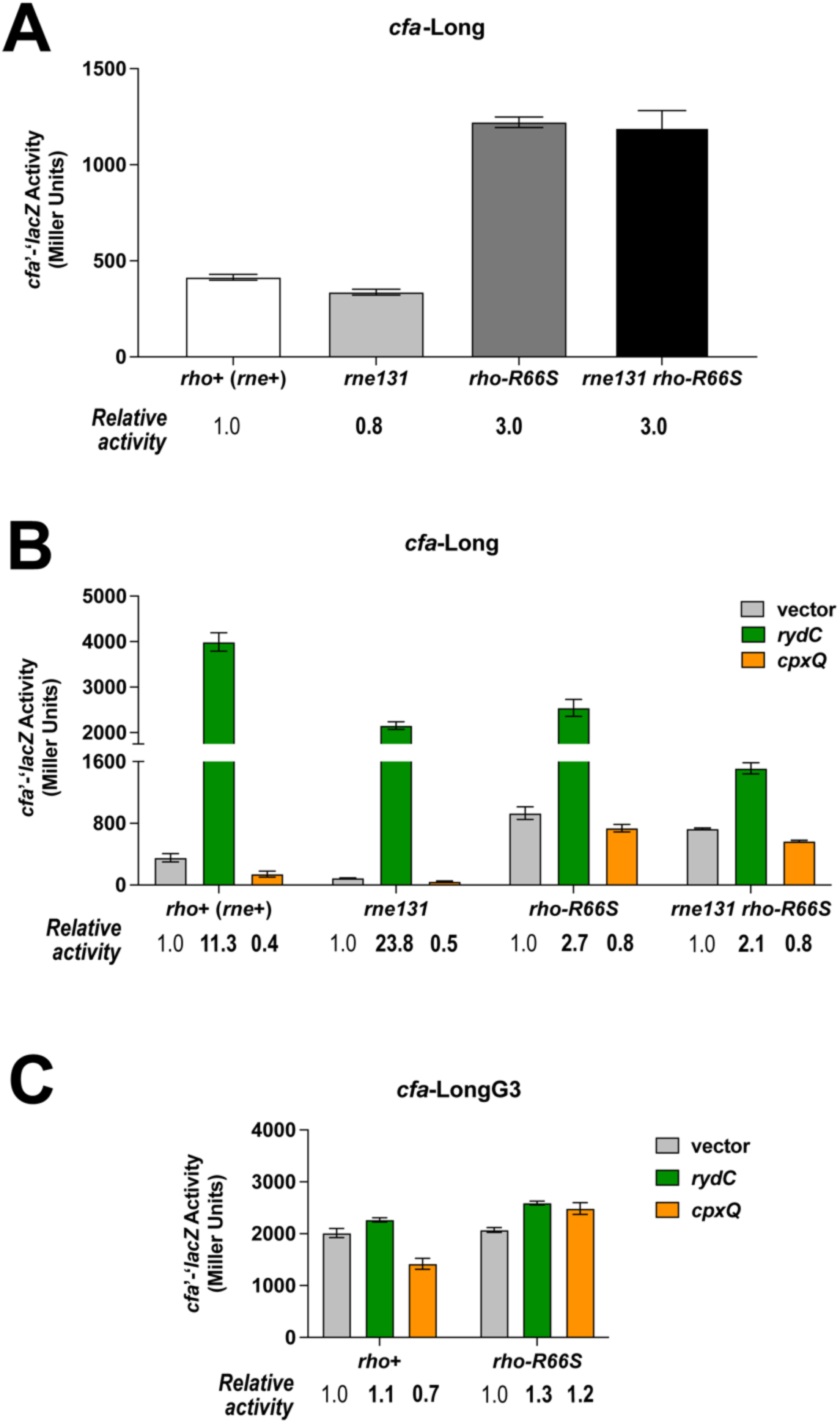
Activating and repressing sRNAs primarily regulate Rho-dependent regulation of *cfa* mRNA. **A**. Activity of the *cfa*-Long translational fusion in wild-type (*rho*^+^ *rne*^+^), *rne131* mutant, *rho*-R66S mutant, and *rne131 rho*-R66S mutant strains. β-galactosidase activity was measured at mid-exponential phase and is expressed in Miller Units. Error bars represent the standard deviations of three biological replicates. **B.** Activity of the *cfa* Long translational fusion in the same strains as in A carrying vector control or plasmids producing RydC or CpxQ. β-galactosidase activity is expressed in Miller Units and the activity was measured at mid-exponential phase after one hour of sRNA induction. Error bars represent the standard deviations of three biological replicates. **C.** Activity of the *cfa*-LongG3 translational fusion in the *rho*^+^ or *rho*-R66S mutant strain background carrying a vector control or plasmid expressing RydC or CpxQ. Error bars represent the standard deviations of three biological replicates.

Previous work demonstrated that *Salmonella cfa* mRNA is cleaved by RNase E at a position adjacent to the RydC binding site (16). Mutation of the cleavage site increased *cfa* mRNA levels and made *cfa* less sensitive to RydC-mediated regulation (12). We made a similar mutation in the *E. coli cfa* 5’ UTR (GG↓A<u>UUA</u>U to GG↓A<u>GGG</u>U, where the arrow indicates the site of cleavage) and showed that the mutation increased *cfa* fusion activity and diminished regulation by RydC and CpxQ (11). Our original interpretation was that both RydC and CpxQ modulate RNase E activity on *cfa* mRNA. However, in light of our current results, we hypothesized that the mutation (where UUA is replaced by GGG, which we will call G3 here) impairs Rho-dependent regulation of *cfa*. To test this, we compared activity of the *cfa*-LongG3 fusion in the *rho^+^* and *rho*-R66S backgrounds carrying an empty vector, RydC-, or CpxQ- producing plasmids (Figure 7C). The G3 mutation strongly increases basal levels of *cfa* expression (compare activity of *cfa-*Long in Figure 7A to *cfa-*LongG3 in 7C) and RydC- and CpxQ-mediated regulation is lost or diminished, respectively (Figure 7C). In the *rho-*R66S background, there is no further increase in the activity of the *cfa*-LongG3 fusion (Figure 7C). These data indeed suggest that the G3 mutation disrupts Rho-dependent regulation of *cfa*, and that this explains changes in G3 fusion activity levels and the loss of RydC- and CpxQ-mediated regulation.

### RydC regulates *cfa* even when RNase E is inactivated

To further probe the role of RNase E in regulation of *cfa* in *E. coli*, we measured *cfa* mRNA levels in *rne*^+^ and *rne131* mutant strains carrying vector control or RydC expression plasmids. In the *rne*^+^ strain with the vector control, *cfa* mRNA was undetectable, and levels increased substantially in the strain producing RydC (Figure 8A). In the *rne131* degradosome mutant, *cfa* mRNA levels were low but detectable and RydC substantially increased these levels (Figure 8A). These results along with the results of reporter fusions (Figure 7B) suggest that the degradosome plays a minor role in *cfa* mRNA turnover and RydC increases *cfa* mRNA levels by a degradosome-independent mechanism.

**Figure 8.**
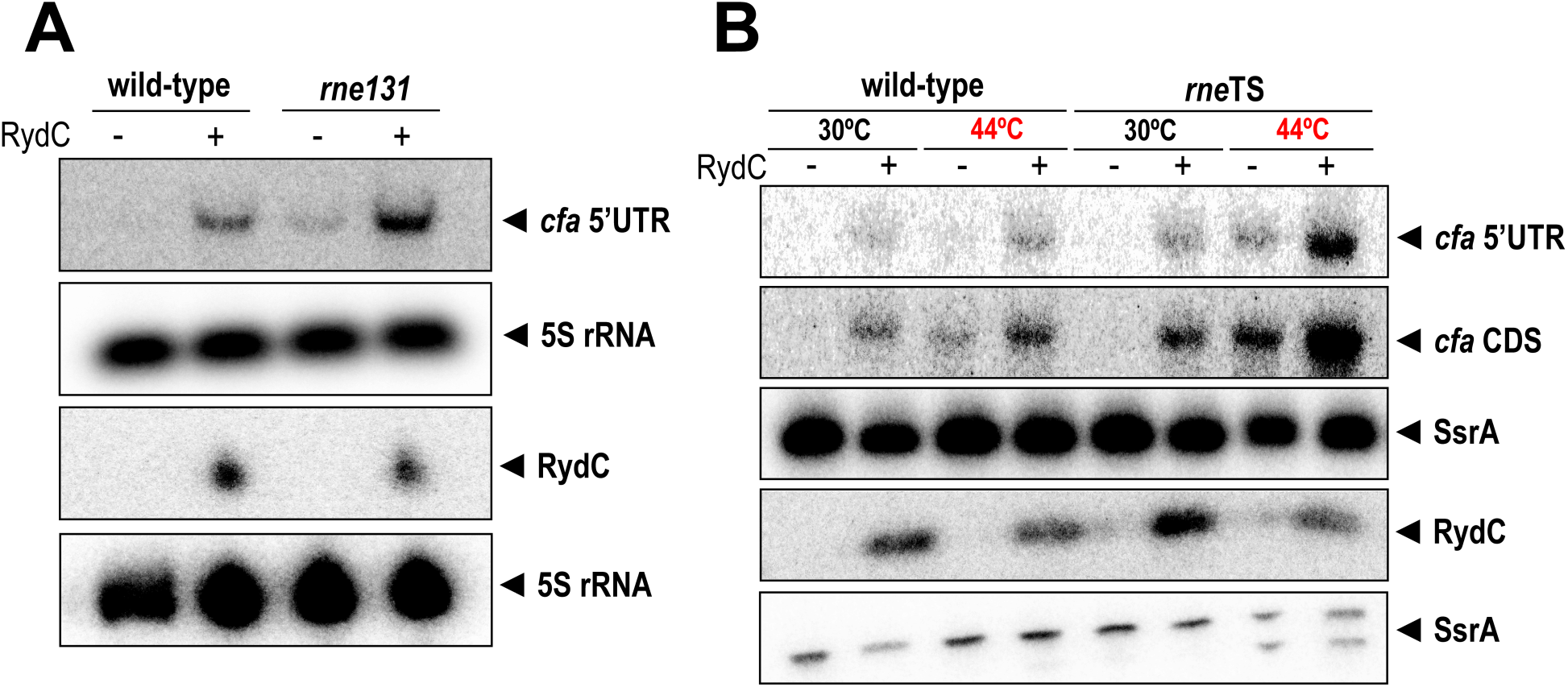
RydC regulates *cfa* even when RNase E is inactivated. **A.** Northern blot analysis was used to examine *cfa* mRNA levels in wild-type (*rne*^+^) and RNase E degradosome mutant (*rne131*) cells in the presence or absence of RydC. The 5S rRNA was used as an RNA loading control. **B.** Northern blot analysis was used to examine *cfa* mRNA levels in wild-type (*rne*^+^) and RNase E temperature-sensitive mutant (*rne*TS) cells at permissive (30°C) and non-permissive (44°C) temperatures in the presence or absence of RydC.

We next compared *cfa* mRNA levels in *rne^+^*and a strain with a temperature-sensitive RNase E (*rne*Ts) (26) each carrying either a vector control or RydC expression plasmid (Figure 8B). In the *rne*^+^ strain grown at both 30°C and 44°C, *cfa* mRNA levels increased when RydC was produced (Figure 8B). In the *rne*Ts strain grown at 30°C, *cfa* mRNA was only detected when RydC was produced. At 44°C, *cfa* mRNA was detectable in the absence of RydC, and levels were further increased when RydC was produced (Figure 8B). If RydC were activating *cfa* primarily by antagonizing RNase E activity, we would not see RydC-dependent accumulation of *cfa* mRNA at the non-permissive temperature where RNase E is inactive. Taken together, our data suggest that while *cfa* mRNA is subject to RNase E-mediated turnover, the sRNAs that regulate *cfa* primarily modulate Rho-dependent transcription termination.

## Discussion

The cyclopropane fatty acid (*cfa*) synthase mRNA is one of a handful of mRNA targets that act as hubs for integration of environmental signals via multiple sRNA regulators. Regulation of *cfa* by multiple sRNAs (11, 12, 16) is particularly intriguing from a mechanistic standpoint because the sRNAs bind at sites distant from the translation initiation region and mediate both positive and negative regulation by translation-independent mechanisms. In this study, we uncovered new details of these mechanisms that expand our understanding of the many varied ways that sRNAs regulate bacterial gene expression. Our data are consistent with a model (Figure 9) where transcription of the long *cfa* isoform is limited by Rho. The long 5’ UTR of *cfa* mRNA contains a *rut* site that includes a CU-rich region required for Rho-dependent regulation. Accessibility of the *rut* site is modulated by the presence of a stem-loop that partially sequesters the CU-rich portion of the *rut* site. CpxQ can repress *cfa* through binding to a site that prevents stem-loop formation, making the *rut* site more accessible and increasing transcription termination (Figure 9, bottom left). RydC can bind to a site downstream of the stem-loop, sequestering a different part of the *rut* site and making it less accessible, leading to decreased transcription termination (Figure 9, bottom right). The sRNAs may also play a minor role in modulating RNase E-mediated *cfa* mRNA turnover, but their major function appears to be control of transcription elongation.

**Figure 9.**
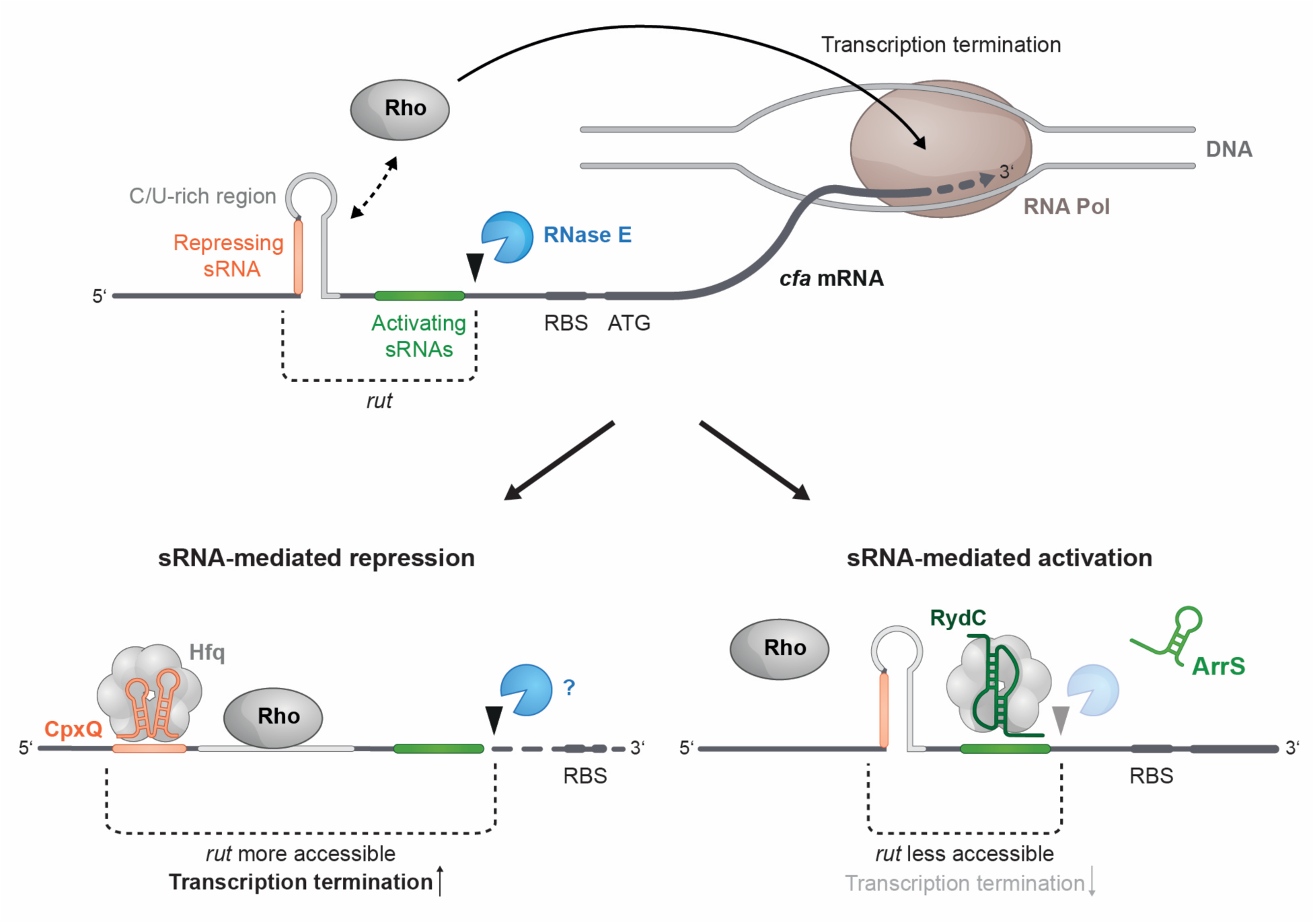
Model for *cfa* regulation by Rho and sRNAs. The model is described in detail in the Discussion.

Regulation of bacterial gene expression at the level of transcription is often thought of as occurring mainly at the level of initiation through the actions of transcription factors that affect RNA polymerase (RNAP) binding to promoter sequences. However, both protein and RNA factors can regulate transcription at the level of elongation through attenuation (increased termination) or antitermination (decreased termination) mechanisms (27). In *E. coli*, 20-30% of transcription termination events are Rho-dependent (28, 29). Because Rho requires a single-stranded binding site unoccupied by ribosomes, regulators or processes that impact translation or RNA structure can influence the efficiency of Rho-dependent termination. Several proteins, including Hfq, have been reported to modulate Rho activity and Rho-dependent termination (30–34). While Hfq is best known as the Sm-like protein involved in sRNA stability and sRNA-mediated regulation, it can also bind directly to Rho to inhibit Rho-dependent termination (32). Small RNAs commonly impact translation and structure of mRNA targets, so it follows that sRNAs have the capacity to control Rho-dependent termination (35). The first translation-dependent mechanism of sRNA control of Rho termination was reported by Bossi and colleagues in 2012 (21). ChiX sRNA represses translation of *chiP* by sequestering the ribosome binding site, which in turn increases the rate of *chiPQ* mRNA turnover (36) and makes a *rut* site in the *chiP* coding sequence more accessible (due to reduced translation), promoting premature Rho-dependent termination (21). Similarly, the sRNA Spot 42 binds to the *galK* ribosome binding site within the polycistronic *galETKM* mRNA. Repressed *galK* translation leads to increased premature Rho-dependent termination within *galK* (37). Our recent work demonstrated that sRNAs including SgrS and RyhB that repress translation of their targets can also promote Rho-dependent termination (38).

Perhaps even more intriguing are a growing number of examples of sRNAs that modulate Rho-dependent termination by apparently translation-independent mechanisms. Long 5’ UTRs are often sites of sRNA action and these sequences are unprotected by ribosomes. Recent work suggested that hundreds of long 5’ UTRs in *E. coli* are targets for Rho-dependent termination (7). Multiple sRNAs activate *rpoS* translation by preventing formation of a translation-inhibitory hairpin in the 5’ UTR (39–42). Recently, these sRNAs were also shown to antagonize premature Rho-dependent termination within the 5’ UTR (7). In *Salmonella*, SraL base pairs with the 5’ UTR of *rho* mRNA to protect it from auto-regulation via Rho-dependent termination (10). For both *rpoS* and *rho*, the mechanism of sRNA-mediated interference with Rho-dependent termination within 5’ UTRs is unclear. To the best of our knowledge, this work is the first to report an sRNA that mediates increased Rho-dependent termination of its mRNA target. Our results demonstrate that the CpxQ binding site is located within the 5’ portion of a stem-loop that partially sequesters sequences important for Rho-dependent termination. We propose that CpxQ binding prevents formation of this structure and increases Rho binding to *cfa* mRNA, promoting premature termination. This is reminiscent of a mechanism used by the RNA binding protein CsrA, which promotes Rho-dependent termination by binding to the *pgaA* mRNA 5’ UTR and remodeling the secondary structure of the 5’ UTR to expose a *rut* site (43).

Classical examples of sRNA-mediated regulation involve direct translational repression through base-pairing interactions that occlude the translation initiation region (44). sRNA-mRNA interactions that occur within a window of approximately 40 nt around the start codon can directly inhibit ribosome binding (45). We now know that sRNAs can also carry out translational regulation of by base pairing at sites that are long distances from translation initiation regions. We and others have described translational repression mechanisms involving sRNA sequestration of translational enhancer sequences (46–48) and ribosome standby sites (49) in long 5’ UTRs. Small RNAs can act as indirect translational repressors by binding upstream (50) or downstream (51) of the translation initiation region and recruiting Hfq to bind at sites overlapping the RBS. Perhaps not surprisingly, regulatory mechanisms that act at steps of gene expression other than translation also involve sRNA-mRNA interactions at sites far from the mRNA translation initiation region. Control of mRNA decay can be modulated by sRNAs binding within 5’ or 3’ UTRs or coding regions (52–54). We can now definitively add transcription elongation to the list of molecular processes that sRNAs can modulate by base pairing at sites distal to the translation initiation region. More work will be required to determine how common sRNA-dependent modulation of transcription elongation is and how both sRNAs and Hfq modulate Rho binding to mRNAs and Rho translocation.

## Materials and Methods

### Bacterial strains, media, and growth conditions

Bacterial strains and plasmids are listed in Table S1. All strains are derivatives of *E. coil* K-12 MG1655. Bacterial complete growth media were purchased from Research Products International (RPI); individual medium components were purchased from Fisher Scientific. Bacteria were cultured in LB broth or on LB agar plates at 37°C unless otherwise specified. Antibiotics and other compounds (from GoldBio unless otherwise specified were used at the following final concentrations: 100 μg/ml ampicillin, 25 μg/ml chloramphenicol, 50 μg/ml kanamycin (Fisher Scientific), 10 μg/ml tetracycline, L-arabinose (Sigma-Aldrich), 0.1 mM isopropyl-β-D-1-thiogalactopyranoside (IPTG), and 40 μg/ml 5-bromo-4-chloro-3-indolyl β-D-galactopyranoside (X-gal).

### Molecular methods

Oligonucleotides (IDT) are listed in Table S2. All reagents and enzymes were purchased from New England Biolabs unless otherwise specified. PCR reactions used Phusion or Q5 High-Fidelity DNA Polymerases. Plasmids were designed using Geneious Prime (v. 2020.1.2) and the NEBuilder Assembly Tool (v. 2.7.1), and plasmids were constructed using the NEBuilder HiFi DNA Assembly Master Mix. Plasmids were verified with restriction enzyme digestion and sequencing.

### Bacterial strain construction

Transcriptional and translational reporter fusions were constructed using λ Red homologous recombination into strain PM1805 and counterselection against *sacB* as previously described (55). gBlock gene fragments were used in construction of *cfa* transcriptional fusions. PCR amplicons or gBlock gene fragments were used to construct *cfa’-’lacZ* and *rho’-’lacZ* translational fusions. All reporter constructs were verified with PCR and sequencing.

A chloramphenicol resistance gene (*cat*) was inserted in the intergenic region between *flgL* and *rne* via λ Red homologous recombination to link the *rne-*3071 (ts) allele to the *cat* gene (56). The *rho-*R66S, *rne-*131, *rne-*3071, Δ*cpxQ,* and Δ*rydC* mutations were moved from donor strains to recipient strains via P1 *vir* transduction (57). Plasmid pCP20 encoding FLP recombinase was used to remove antibiotic resistance cassettes from Δ*cpxQ*::kan and Δ*rydC*::kan strains as needed (56).

### β-galactosidase assays

Bacterial strains harboring *lacZ* reporter fusions were cultured at 37°C for 16 hours in TB (tryptone broth) medium with arabinose with or without ampicillin. After 16 hours, bacterial strains were subcultured 1:100 in the same media and grown to early or mid-exponential phase. If sRNA induction was not required, cultures were harvested and assayed at mid-log phase. If sRNA induction was required, 0.1 mM IPTG was added at early exponential phase and cells were cultured at 37°C for 1 hour more before harvesting and assaying at mid-log phase. β- galactosidase assays were carried out with chloroform-permeabilized cells as previously described (57) and data are expressed in Miller Units.

### RNA extraction

Total RNA was extracted from cells using the hot phenol method as previously described (58) with the following adjustments. 1 ml of culture was mixed with 1 ml of 65°C lysis solution containing SDS, EDTA, sodium acetate, and UltraPure Phenol:Water (3.75:1, v/v) (Invitrogen), and the mixture was shaken at 65°C/1400 rpm for 15 minutes. Samples were centrifuged at room temperature/21,300 x g for 10 minutes. The aqueous layer was transferred to a 5PRIME Phase Lock Gel Heavy wax tube (Quantabio) and then extracted with one volume of Phenol:Chloroform:Isoamyl alcohol (25:24:1) (Ambion). The aqueous layer was transferred to a tube containing 1.3 ml of absolute ethanol and placed at −80°C. The next day, the sample was centrifuged at 4°C/14,000 x g for 30 minutes. The pellet was allowed to dry before resuspension in 10-40 μl of RNase-free water. RNA samples were stored at −80°C. RNA samples were treated with TURBO DNase (Invitrogen) according to the manufacturer’s instructions. After DNase digestion, samples were purified using phenol-chloroform extraction. Two volumes of RNase-free water and 0.2 volumes of sodium acetate were added to each reaction, an equivalent volume of acid phenol:chloroform (pH 4.5) (Ambion) was added, and each sample was transferred to Phase Lock Gel Heavy wax tube. RNA was precipitated, then resuspended as described above. RNA concentrations were measured using the Qubit RNA Broad Range (BR) Assay kit.

### Northern blot analysis

DNA probes (IDT) were radiolabled at the 5’ end with fresh [γ-^32^P] ATP (Perkin Elmer) using the KinaseMax Kit (Invitrogen) according to the manufacturer’s instructions. Radiolabeled probes were purified using ProbeQuant G-50 Micro Columns (Cytiva) and were stored at −80°C until use. Nylon membranes were stripped and re-probed as previously described (11).

For analysis of *cfa* mRNA levels, 20 μg RNA samples were denatured at 65°C for 15 minutes in loading buffer (containing MOPS buffer, formaldehyde, and formamide), separated on a 0.8% agarose gel for electrophoresis at 80 V for 2 hours in 1X MOPS buffer, and transferred to a BrightStar-Plus Positively Charged Nylon Membrane (Invitrogen) via capillary transfer overnight. Transferred RNAs were UV crosslinked to the membrane, and the membrane was prehybridized with 10 ml ULTRAhyb Ultrasensitive Hybridization Buffer (Invitrogen) at 42°C for 45 minutes. Blots were allowed to hybridize with radiolabeled probes overnight at 42°C and were then washed once with 2X SSC/0.1% SDS, once with 0.1X SSC/0.1% SDS, and once with 0.1X SSC for 15 minutes each. For signal detection, blots were exposed to a phosphorimager screen and the results were visualized using a Typhoon FLA 9500 Phosphorimager (GE).

For analysis of sRNA levels, 10 μg RNA samples were prepared and blotted as previously described (11) with the following modifications. Electroblot transfer was run on ice at 250 mA (constant) for 4 hours with 0.5X TBE buffer. After hybridization, blots were washed once with 2X SSC/0.1% SDS, once with 0.1X SSC/0.1% SDS, and once with 0.1X SSC for 15 minutes at 42°C. For signal detection, blots were exposed to a phosphorimager screen and the results were visualized using a Typhoon FLA 9500 Phosphorimager (GE).

### Reverse transcription and quantitative PCR (RT-qPCR)

RNA extracts were treated with TURBO DNase (Invitrogen) as described above. Samples were verified to be DNA-free using 40-cycle PCR reactions with each PCR primer pair. The RNA concentration was quantified using the Qubit RNA Broad Range (BR) kit. RT-qPCR reactions (10 μl) contained 250 ng RNA, forward primer, reverse primer, probe, and Luna Probe One-Step RT-qPCR 4X Mix with UDG (NEB) according to the manufacturer’s instructions. Control reactions contained Luna Probe One-Step RT-qPCR 4X Mix with UDG that was first heated at 95°C for 5 minutes to inactivate RT. RT-qPCR reactions were run on a Bio-Rad CFX Connect Real-Time System with Bio-Rad CFX Manager (Version 3.1) software in 0.2 ml 96-well PCR plates (dot scientific inc.) sealed with transparent Microseal B Adhesive Sealer (Bio-Rad). Data were analyzed using the comparative Ct (2^-ΔΔCt^) method.

## Acknowledgements

We thank Asma Hatoum-Aslan, Jim Slauch, and members of the Slauch lab for thoughtful discussions and advice. We also thank past and present members of the Vanderpool lab for strains, phage, plasmids, and advice. Special thanks to Rachel Mooney and Robert Landick for strains, plasmids and advice. We are grateful to Dr. Sandy Westermann of Scigraphix for graphic design. This work was supported by the National Institutes of Health R35 grant (R35 GM139557, to C.K.V.) and the GEMS Biology Integration Institute, funded by the National Science Foundation DBI Biology Integration Institutes Program, award no. 2022049 (K.R.F, A.K.B., C.K.V.).

## Notes

### Competing Interest Statement

The authors have declared no competing interest.

### Summary of Updates

New data clarifying the mechanism of regulation of cfa mRNA by RydC and CpxQ sRNAs were added (new Fig. 6). The author who completed this work has been added (Andrew K. Buechler).

## References

1. Carvalho Barbosa C, Calhoun SH, Wieden HJ. 2020. Non-coding RNAs: what are we missing? Biochem Cell Biol 98:23–30.

2. Adams PP, Storz G. 2020. Prevalence of small base-pairing RNAs derived from diverse genomic loci. Biochim Biophys Acta Gene Regul Mech 1863:194524.

3. Hor J, Matera G, Vogel J, Gottesman S, Storz G. 2020. Trans-Acting Small RNAs and Their Effects on Gene Expression in *Escherichia coli* and *Salmonella enterica*. EcoSal Plus 9.

4. Jørgensen MG, Pettersen JS, Kallipolitis BH. 2020. sRNA-mediated control in bacteria: An increasing diversity of regulatory mechanisms. Biochim Biophys Acta Gene Regul Mech 1863:194504.

5. Papenfort K, Vanderpool CK. 2015. Target activation by regulatory RNAs in bacteria. FEMS Microbiol Rev 39:362–78.

6. Bossi L, Figueroa-Bossi N, Bouloc P, Boudvillain M. 2020. Regulatory interplay between small RNAs and transcription termination factor Rho. Biochim Biophys Acta Gene Regul Mech 1863:194546.

7. Sedlyarova N, Shamovsky I, Bharati BK, Epshtein V, Chen J, Gottesman S, Schroeder R, Nudler E. 2016. sRNA-Mediated Control of Transcription Termination in E. coli. Cell 167:111–121 e13.

8. Boudvillain M, Figueroa-Bossi N, Bossi L. 2013. Terminator still moving forward: expanding roles for Rho factor. Curr Opin Microbiol 16:118–24.

9. Kung H, Bekesi E, Guterman SK, Gray JE, Traub L, Calhoun DH. 1984. Autoregulation of the *rho* gene of *Escherichia coli* K-12. Mol Gen Genet 193:210–3.

10. Silva IJ, Barahona S, Eyraud A, Lalaouna D, Figueroa-Bossi N, Masse E, Arraiano CM. 2019. SraL sRNA interaction regulates the terminator by preventing premature transcription termination of rho mRNA. Proc Natl Acad Sci U S A 116:3042–3051.

11. Bianco CM, Fröhlich KS, Vanderpool CK. 2019. Bacterial Cyclopropane Fatty Acid Synthase mRNA Is Targeted by Activating and Repressing Small RNAs. J Bacteriol 201.

12. Fröhlich KS, Papenfort K, Fekete A, Vogel J. 2013. A small RNA activates CFA synthase by isoform-specific mRNA stabilization. EMBO J 32:2963–79.

13. Grogan DW, Cronan JE, Jr. 1997. Cyclopropane ring formation in membrane lipids of bacteria. Microbiol Mol Biol Rev 61:429–41.

14. Shabala L, Ross T. 2008. Cyclopropane fatty acids improve *Escherichia coli* survival in acidified minimal media by reducing membrane permeability to H+ and enhanced ability to extrude H+. Res Microbiol 159:458–61.

15. Chao Y, Vogel J. 2016. A 3’ UTR-Derived Small RNA Provides the Regulatory Noncoding Arm of the Inner Membrane Stress Response. Mol Cell 61:352–363.

16. Chao Y, Li L, Girodat D, Forstner KU, Said N, Corcoran C, Smiga M, Papenfort K, Reinhardt R, Wieden HJ, Luisi BF, Vogel J. 2017. In Vivo Cleavage Map Illuminates the Central Role of RNase E in Coding and Non-coding RNA Pathways. Mol Cell 65:39–51.

17. Adams PP, Baniulyte G, Esnault C, Chegireddy K, Singh N, Monge M, Dale RK, Storz G, Wade JT. 2021. Regulatory roles of *Escherichia coli* 5’ UTR and ORF-internal RNAs detected by 3’ end mapping. Elife 10.

18. Wang AY, Cronan JE, Jr. 1994. The growth phase-dependent synthesis of cyclopropane fatty acids in *Escherichia coli* is the result of an RpoS(KatF)-dependent promoter plus enzyme instability. Mol Microbiol 11:1009–17.

19. Martinez A, Opperman T, Richardson JP. 1996. Mutational analysis and secondary structure model of the RNP1-like sequence motif of transcription termination factor Rho. J Mol Biol 257:895–908.

20. Matsumoto Y, Shigesada K, Hirano M, Imai M. 1986. Autogenous regulation of the gene for transcription termination factor *rho* in *Escherichia coli*: localization and function of its attenuators. J Bacteriol 166:945–58.

21. Bossi L, Schwartz A, Guillemardet B, Boudvillain M, Figueroa-Bossi N. 2012. A role for Rho-dependent polarity in gene regulation by a noncoding small RNA. Genes Dev 26:1864–73.

22. Lopez PJ, Marchand I, Joyce SA, Dreyfus M. 1999. The C-terminal half of RNase E, which organizes the *Escherichia coli* degradosome, participates in mRNA degradation but not rRNA processing *in vivo*. Mol Microbiol 33:188–99.

23. Bobrovskyy M, Vanderpool CK. 2016. Diverse mechanisms of post-transcriptional repression by the small RNA regulator of glucose-phosphate stress. Mol Microbiol 99:254–73.

24. Caron MP, Lafontaine DA, Masse E. 2010. Small RNA-mediated regulation at the level of transcript stability. RNA Biol 7:140–4.

25. Rice JB, Balasubramanian D, Vanderpool CK. 2012. Small RNA binding-site multiplicity involved in translational regulation of a polycistronic mRNA. Proc Natl Acad Sci U S A 109:E2691–8.

26. McDowall KJ, Hernandez RG, Lin-Chao S, Cohen SN. 1993. The *ams-1* and *rne-*3071 temperature-sensitive mutations in the ams gene are in close proximity to each other and cause substitutions within a domain that resembles a product of the *Escherichia coli* mre locus. J Bacteriol 175:4245–9.

27. Turnbough CL, Jr. 2019. Regulation of Bacterial Gene Expression by Transcription Attenuation. Microbiol Mol Biol Rev 83.

28. Peters JM, Mooney RA, Grass JA, Jessen ED, Tran F, Landick R. 2012. Rho and NusG suppress pervasive antisense transcription in *Escherichia coli*. Genes Dev 26:2621–33.

29. Peters JM, Mooney RA, Kuan PF, Rowland JL, Keles S, Landick R. 2009. Rho directs widespread termination of intragenic and stable RNA transcription. Proc Natl Acad Sci U S A 106:15406–11.

30. Gutierrez P, Kozlov G, Gabrielli L, Elias D, Osborne MJ, Gallouzi IE, Gehring K. 2007. Solution structure of YaeO, a Rho-specific inhibitor of transcription termination. J Biol Chem 282:23348–53.

31. Pal K, Yadav M, Jain S, Ghosh B, Sen R, Sen U. 2019. *Vibrio cholerae* YaeO is a Structural Homologue of RNA Chaperone Hfq that Inhibits Rho-dependent Transcription Termination by Dissociating its Hexameric State. J Mol Biol 431:4749–4766.

32. Rabhi M, Espeli O, Schwartz A, Cayrol B, Rahmouni AR, Arluison V, Boudvillain M. 2011. The Sm-like RNA chaperone Hfq mediates transcription antitermination at Rho-dependent terminators. EMBO J 30:2805–16.

33. Said N, Finazzo M, Hilal T, Wang B, Selinger TL, Gjorgjevikj D, Artsimovitch I, Wahl MC. 2024. Sm-like protein Rof inhibits transcription termination factor *rho* by binding site obstruction and conformational insulation. Nat Comm 15:3186.

34. Wang B, Pei H, Lu Z, Xu Y, Han S, Jia Z, Zheng J. 2022. YihE is a novel binding partner of Rho and regulates Rho-dependent transcription termination in the Cpx stress response. iScience 25:105483.

35. Chen J, Morita T, Gottesman S. 2019. Regulation of Transcription Termination of Small RNAs and by Small RNAs: Molecular Mechanisms and Biological Functions. Front Cell Infect Microbiol 9:201.

36. Figueroa-Bossi N, Valentini M, Malleret L, Fiorini F, Bossi L. 2009. Caught at its own game: regulatory small RNA inactivated by an inducible transcript mimicking its target. Genes Dev 23:2004–15.

37. Wang X, Ji SC, Jeon HJ, Lee Y, Lim HM. 2015. Two-level inhibition of *galK* expression by Spot 42: Degradation of mRNA mK2 and enhanced transcription termination before the *galK* gene. Proc Natl Acad Sci U S A 112:7581–6.

38. Reyer MA, Chennakesavalu S, Heideman EM, Ma X, Bujnowska M, Hong L, Dinner AR, Vanderpool CK, Fei J. 2021. Kinetic modeling reveals additional regulation at co-transcriptional level by post-transcriptional sRNA regulators. Cell Rep 36:109764.

39. Battesti A, Majdalani N, Gottesman S. 2011. The RpoS-mediated general stress response in *Escherichia coli*. Annu Rev Microbiol 65:189–213.

40. Majdalani N, Cunning C, Sledjeski D, Elliott T, Gottesman S. 1998. DsrA RNA regulates translation of RpoS message by an anti-antisense mechanism, independent of its action as an antisilencer of transcription. Proc Natl Acad Sci U S A 95:12462–7.

41. Majdalani N, Hernandez D, Gottesman S. 2002. Regulation and mode of action of the second small RNA activator of RpoS translation, RprA. Mol Microbiol 46:813–26.

42. Mandin P, Gottesman S. 2010. Integrating anaerobic/aerobic sensing and the general stress response through the ArcZ small RNA. EMBO J 29:3094–107.

43. Figueroa-Bossi N, Schwartz A, Guillemardet B, D’Heygere F, Bossi L, Boudvillain M. 2014. RNA remodeling by bacterial global regulator CsrA promotes Rho-dependent transcription termination. Genes Dev 28:1239–51.

44. De Lay N, Schu DJ, Gottesman S. 2013. Bacterial small RNA-based negative regulation: Hfq and its accomplices. J Biol Chem 288:7996–8003.

45. Bouvier M, Sharma CM, Mika F, Nierhaus KH, Vogel J. 2008. Small RNA binding to 5’ mRNA coding region inhibits translational initiation. Mol Cell 32:827–37.

46. Azam MS, Vanderpool CK. 2020. Translation inhibition from a distance: The small RNA SgrS silences a ribosomal protein S1-dependent enhancer. Mol Microbiol 114:391–408.

47. Sharma CM, Darfeuille F, Plantinga TH, Vogel J. 2007. A small RNA regulates multiple ABC transporter mRNAs by targeting C/A-rich elements inside and upstream of ribosome-binding sites. Genes Dev 21:2804–17.

48. Yang Q, Figueroa-Bossi N, Bossi L. 2014. Translation enhancing ACA motifs and their silencing by a bacterial small regulatory RNA. PLoS Genet 10:e1004026.

49. Darfeuille F, Unoson C, Vogel J, Wagner EG. 2007. An antisense RNA inhibits translation by competing with standby ribosomes. Mol Cell 26:381–92.

50. Desnoyers G, Masse E. 2012. Noncanonical repression of translation initiation through small RNA recruitment of the RNA chaperone Hfq. Genes Dev 26:726–39.

51. Azam MS, Vanderpool CK. 2018. Translational regulation by bacterial small RNAs via an unusual Hfq-dependent mechanism. Nucleic Acids Res 46:2585–2599.

52. Fröhlich KS, Papenfort K, Berger AA, Vogel J. 2012. A conserved RpoS-dependent small RNA controls the synthesis of major porin OmpD. Nucleic Acids Res 40:3623–40.

53. Papenfort K, Sun Y, Miyakoshi M, Vanderpool CK, Vogel J. 2013. Small RNA-mediated activation of sugar phosphatase mRNA regulates glucose homeostasis. Cell 153:426–37.

54. Pfeiffer V, Papenfort K, Lucchini S, Hinton JC, Vogel J. 2009. Coding sequence targeting by MicC RNA reveals bacterial mRNA silencing downstream of translational initiation. Nat Struct Mol Biol 16:840–6.

55. Mandin P, Gottesman S. 2009. A genetic approach for finding small RNAs regulators of genes of interest identifies RybC as regulating the DpiA/DpiB two-component system. Mol Microbiol 72:551–65.

56. Datsenko KA, Wanner BL. 2000. One-step inactivation of chromosomal genes in *Escherichia coli* K-12 using PCR products. Proc Natl Acad Sci U S A 97:6640–5.

57. Miller JH. 1972. Experiments in Molecular Genetics. Cold Spring Harbor Laboratory Press.

58. Aiba H, Adhya S, de Crombrugghe B. 1981. Evidence for two functional *gal* promoters in intact *Escherichia coli* cells. J Biol Chem 256:11905–10.

