## Supplemental Figs and Tables for "Small RNAs positively and negatively control transcription elongation through modulation of Rho utilization site accessibility"

**Supplementary Material**

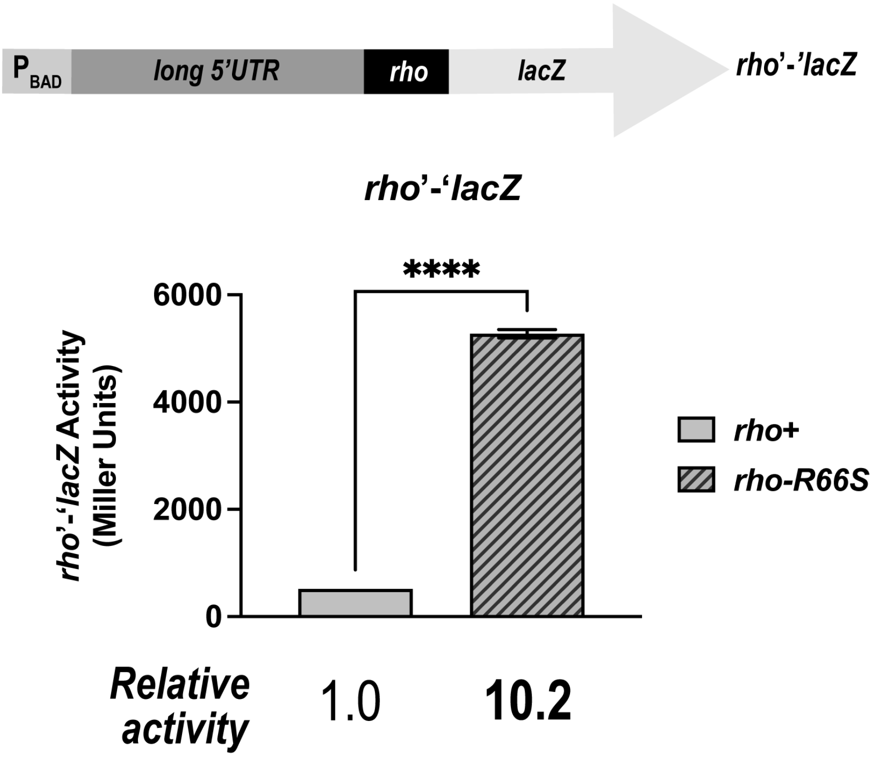


**Figure S1. A *rho* reporter fusion responds as expected to the *rho*-R66S allele.** A *rho*'-'*lacZ* translational fusion contains the entire 5' UTR of *rho* mRNA and the first 10 codons of the *rho* coding sequence (top). Transcription of the fusion was controlled with an arabinose-inducible promoter ($P_{BAD}$). β-galactosidase activity of the *rho*'-'*lacZ* fusion was tested in *rho*+ and *rho*-R66S mutant backgrounds (bottom). β-galactosidase activity is expressed in Miller Units and the activity was measured at mid-exponential phase. Error bars represent the standard deviations of three biological replicates and statistical significance was determined using a two-tailed Welch's t-test (****$P < 0.0001$).

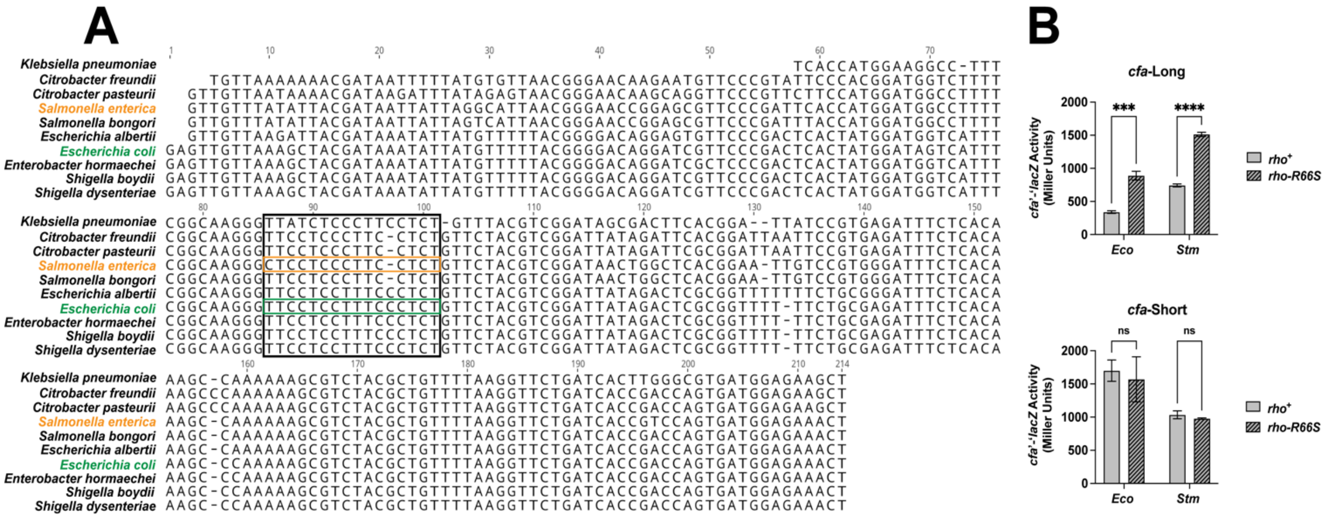


**Figure S2. The long 5' UTR of *cfa* mRNA from *Salmonella enterica* is required for premature Rho-dependent transcription termination. A.** Nucleotide sequence alignment of *cfa* mRNA 5' UTR sequences from ten different enterobacterial species. The larger boxed region contains the portion of each sequence aligned with the *E. coli* CU-rich region (green box). The orange box highlights the *S. enterica cfa* CU-rich sequence. The alignment was generated using the Clustal Omega program (version 1.2.3) through Geneious Prime software (version 2023.2.1). **B.** β-galactosidase activity of *E. coli* (*Eco*) and *S. enterica* (*Stm*) *cfa*-Long and *cfa*-Short fusions was measured at mid-exponential phase. Error bars represent the standard deviations of three biological replicates and statistical significance was determined using two-tailed Welch's t-tests (****P < 0.001).

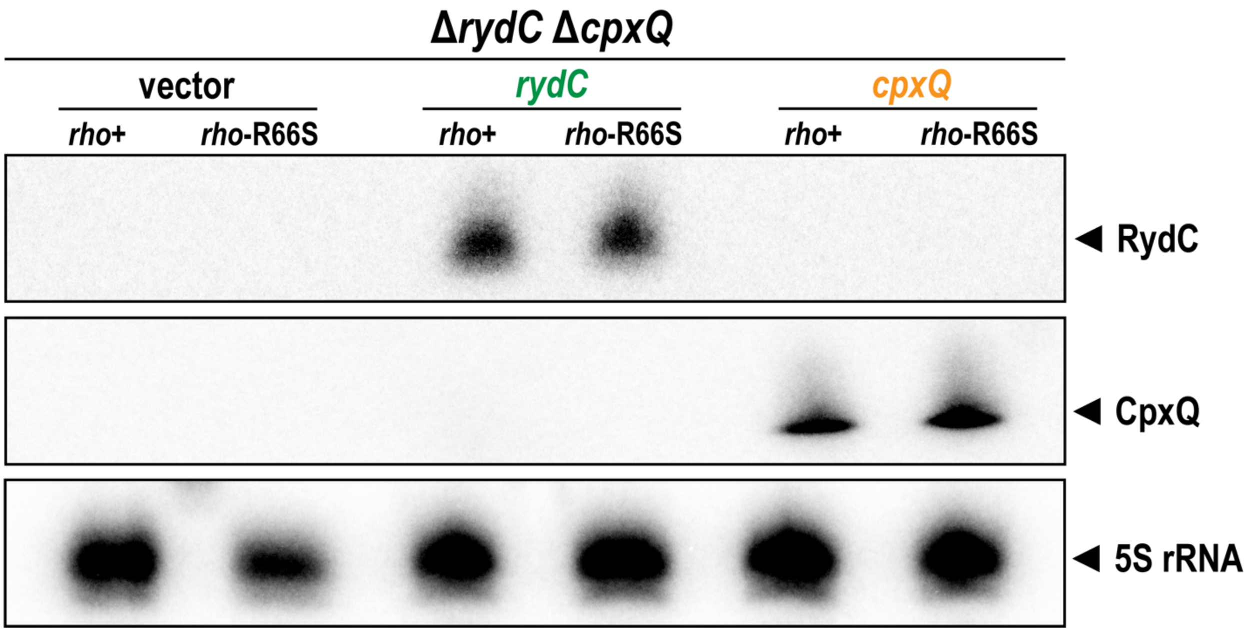


**Figure S3. RydC and CpxQ are expressed at similar levels in wild-type and *rho* mutant strains.** Northern blot analysis to examine the levels of RydC and CpxQ sRNAs in the presence of a vector control or RydC- or CpxQ-producing plasmids in wild-type (Δ*rydC* Δ*cpxQ* *rho*+) and *rho* mutant (Δ*rydC* Δ*cpxQ* *rho*-R66S) cells. The 5S rRNA was used as an RNA loading control.

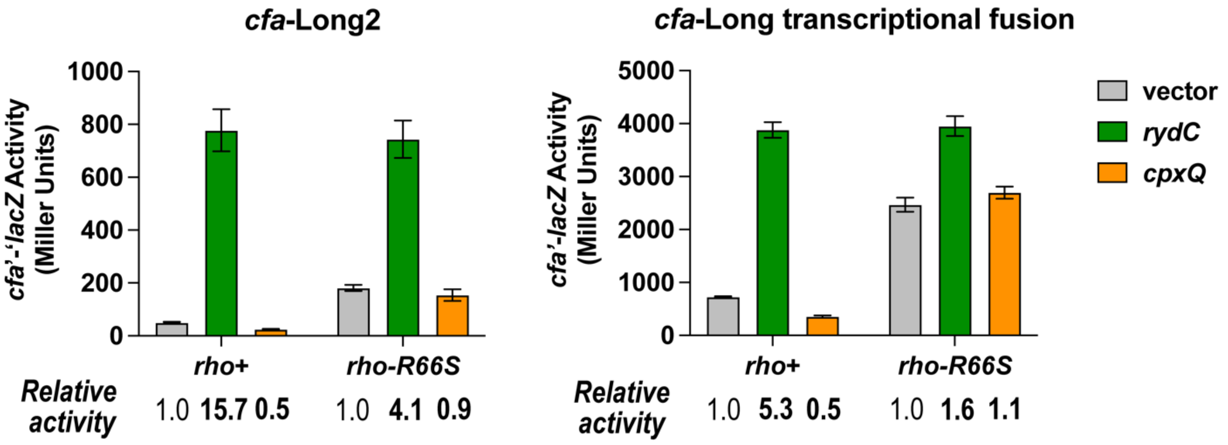


**Figure S4. sRNA-dependent regulation of the *cfa*-Long2 translational fusion and *cfa*-Long transcriptional fusion.** β-galactosidase activity of the *cfa*-Long2 translational fusion and the *cfa*-Long transcriptional fusion was measured at mid-exponential phase in *rho*+ and *rho*-R66S mutant backgrounds in presence of a vector control or RydC- or CpxQ-producing plasmids. Error bars represent the standard deviations of three biological replicates.

**Table S1. Plasmids and strains used in this study.**

| Plasmid | Vector | Genotype | Source or Reference |
|---|---|---|---|
| pBRCS12 | pHDB3 | vector control | Wadler et al., 2009 |
| pCB4 | pBRCS12 | $P_{lac}$-*arrS* | Bianco 2019 |
| pCB5 | pBRCS12 | $P_{lac}$-*cpxQ* | Bianco 2019 |
| pCB7 | pBRCS12 | $P_{lac}$-*rydC* | Bianco 2019 |
| pCB11 | pBRCS12 | $P_{lac}$-*oxyS* | Bianco 2019 |
| pCB12 | pBRCS12 | $P_{lac}$-*gcvB* | Bianco 2019 |
| pAKB001 | pBRCS12 | $P_{lac}$-*cpxQ_C10G_C11G* | This study |
| pAKB002 | pBRCS12 | $P_{lac}$-*cpxQ_G9U* | This study |
| **Strain** | **Parent** | **Genotype** | **Source or Reference** |
| DJ480 | MG1655 | Δ*lac* X74 | D. Jin, NCI |
| EM1277 | DJ480 (EM1055) | Δ*lac* X74 *rne-3071* (ts) *zce-726*::Tn10 | Masse, Escorcia, and Gottesman 2003 |
| PM1805 | PM1205 | MG1655 *mal*::*lacI*$^{Q}$ Δ*araBAD araC*+ *lacI*'::$P_{BAD}$-*cat-sacB-lacZ* miniλ-tet$^{R}$ | Lee and Gottesman 2016 |
| AK27 | PM1205 | $P_{BAD}$-*cfa*'-'*lacZ*-Long | King 2019 |
| AK28 | PM1205 | $P_{BAD}$-*cfa*'-'*lacZ*-Short | King 2019 |
| CB839 | PM1205 | $P_{BAD}$-*cfa*'-'*lacZ*-LongG3 | Bianco 2019 |
| CB1118 | CB839 | $P_{BAD}$-*cfa*'-'*lacZ*-LongG3 *rho*-R66S | This study |
| CB1089 | DJ480 | Δ*lac* X74 *rho*-R66S | This study |
| CB1097 | AK28 | $P_{BAD}$-*cfa*'-'*lacZ*-Short *rho*-R66S | This study |
| CB1098 | AK27 | $P_{BAD}$-*cfa*'-'*lacZ*-Long *rho*-R66S | This study |
| KF5 | PM1805 | $P_{BAD}$-*cfa*'-'*lacZ*-Long2 | This study |
| KF6 | PM1805 | $P_{BAD}$-*cfa*'-'*lacZ*-Short2 | This study |
| KF16 | KF5 | $P_{BAD}$-*cfa*'-'*lacZ*-Long2 *rho*-R66S | This study |
| KF17 | KF6 | $P_{BAD}$-*cfa*'-'*lacZ*-Short2 *rho*-R66S | This study |
| KF43 | PM1805 | $P_{BAD}$-*rho*'-'*lacZ* | This study |
| KF61 | KF43 | $P_{BAD}$-*rho*'-'*lacZ* *rho*-R66S | This study |
| KF88 | PM1805 | $P_{BAD}$-*cfa*'-'*lacZ*-Long G fusion | This study |
| KF92 | PM1805 | $P_{BAD}$-*cfa*'-'*lacZ*-Long GGG fusion | This study |
| KF94 | PM1805 | $P_{BAD}$-*cfa*'-'*lacZ*-Long GGGGGG fusion | This study |

| | | | |
|---|---|---|---|
| KF100 | KF88 | $P_{BAD}$-*cfa*'-'*lacZ*-Long G fusion *rho*-R66S | This study |
| KF102 | KF92 | $P_{BAD}$-*cfa*'-'*lacZ*-Long GGG fusion *rho*-R66S | This study |
| KF103 | KF94 | $P_{BAD}$-*cfa*'-'*lacZ*-Long GGGGGG fusion *rho*-R66S | This study |
| KF130 | KF128 | Δ*cpxQ* Δ*rydC* | This study |
| KF145 | PM1805 | $P_{BAD}$-*cfa*'-'*lacZ*-Long (*S. enterica*) | This study |
| KF147 | PM1805 | $P_{BAD}$-*cfa*'-'*lacZ*-Short (*S. enterica*) | This study |
| KF149 | KF145 | $P_{BAD}$-*cfa*'-'*lacZ*-Long (*S. enterica*) *rho*-R66S | This study |
| KF150 | KF147 | $P_{BAD}$-*cfa*'-'*lacZ*-Short (*S. enterica*) *rho*-R66S | This study |
| KF152 | KF130 | Δ*cpxQ* Δ*rydC* *rho*-R66S | This study |
| KF181 | AK27 | $P_{BAD}$-*cfa*'-'*lacZ*-Long *rne131* | This study |
| KF184 | KF181 | $P_{BAD}$-*cfa*'-'*lacZ*-Long *rne131* *rho*-R66S | This study |
| KF217 | DJ480 | Δ*lac* X74 *rne131* | This study |
| KF264 | PM1805 | $P_{BAD}$-*cfa*'-*lacZ*-Long | This study |
| KF265 | PM1805 | $P_{BAD}$-*cfa*'-*lacZ*-Short | This study |
| KF266 | KF264 | $P_{BAD}$-*cfa*'-*lacZ*-Long *rho*-R66S | This study |
| KF267 | KF265 | $P_{BAD}$-*cfa*'-*lacZ*-Short *rho*-R66S | This study |
| KF279 | EM1277 | Δ*lac* X74 *rne-3071* (ts) $Cm^R$ *zce-726*::Tn10 | This study |
| KF308 | KF279 | Δ*lac* X74 *rne-3071* (ts) $Cm^R$ *zce-726*::Tn10 *rho*-R66S | This study |
| KF290 | PM1805 | $P_{BAD}$-*cfa*'-*lacZ*-Long +58 | This study |
| KF291 | PM1805 | $P_{BAD}$-*cfa*'-*lacZ*-Long +78 | This study |
| KF292 | PM1805 | $P_{BAD}$-*cfa*'-*lacZ*-Long +98 | This study |
| KF293 | PM1805 | $P_{BAD}$-*cfa*'-*lacZ*-Long +118 | This study |
| KF294 | PM1805 | $P_{BAD}$-*cfa*'-*lacZ*-Long +138 | This study |
| KF295 | PM1805 | $P_{BAD}$-*cfa*'-*lacZ*-Long +158 | This study |
| KF300 | KF290 | $P_{BAD}$-*cfa*'-*lacZ*-Long +58 *rho*-R66S | This study |
| KF301 | KF291 | $P_{BAD}$-*cfa*'-*lacZ*-Long +78 *rho*-R66S | This study |
| KF302 | KF292 | $P_{BAD}$-*cfa*'-*lacZ*-Long +98 *rho*-R66S | This study |

| | | | |
|---|---|---|---|
| KF303 | KF293 | $P_{BAD}$-*cfa*'-*lacZ*-Long +118 *rho*-R66S | This study |
| KF304 | KF294 | $P_{BAD}$-*cfa*'-*lacZ*-Long +138 *rho*-R66S | This study |
| KF305 | KF295 | $P_{BAD}$-*cfa*'-*lacZ*-Long +158 *rho*-R66S | This study |
| AKB078 | KF264 | $P_{BAD}$-*cfa*'-'*lacZ*-Long G78C/G79C (WS) | This study |
| AKB079 | KF264 | $P_{BAD}$-*cfa*'-'*lacZ*-Long G78C/G79C (WS) *rho*-R66S | This study |
| AKB080 | KF264 | $P_{BAD}$-*cfa*'-'*lacZ*-Long C80A (SS) | This study |
| AKB081 | KF264 | $P_{BAD}$-*cfa*'-'*lacZ*-Long C80A (SS) *rho*-R66S | This study |

**Table 2. Oligonucleotides used in this study.**

| Oligo | Description | Sequence (5' to 3') |
|---|---|---|
| *cfa*-Long2 Forward | Forward primer for *cfa*'-'*lacZ*-Long2 fusion construction | ACCTGACGCTTTTTATCGCAACTCTCTACTGTTT CTCCATGAGTTGTTAAAGCTACGATA |
| *cfa*-Short2 Forward | Forward primer for *cfa*'-'*lacZ*-Short2 fusion construction | ACCTGACGCTTTTTATCGCAACTCTCTACTGTTTCTC CATAAGGTTCTGATCACCGACCA |
| *cfa*-2 Reverse | Reverse primer for *cfa*'-'*lacZ*-Long2 and *cfa*'-'*lacZ*-Short2 fusion construction | TAACGCCAGGGTTTTCCCAGTCACGACGTTGTAAA ACGACGACGCCCACCACGCTTACGT |
| *cfa*-Long G mutant gBlock | gBlock gene fragment for *cfa*'-'*lacZ*-Long G fusion construction | TCGCAACTCTCTACTGTTTCTCCATGAGTTGTTAAAG CTACGATAAATATTATGTTTTTACGGGGACAGGATCG TTCCCGACTCACTATGGATAGTCATTTCGGCAAGGGT TCCTCCTTTCCCTGTGTTCTACGTCGGATTATAGACT CGCGGTTTTTTCTGCGAGATTTCTCACAAAGCCCAAA AAGCGTCTACGCTGTTTTAAGGTTCTGATCACCGACC AGTGATGGAGAAACTATGAGTTCATCGTGTATAGAA GAAGTCAGTGTACCGGATGACAACTGGTACCGTATC GCCAACGAATTACTTAGCCGTGCCGGTATAGCCATT AACGGTTCTGCCCCGGTCGTTTTACAACGTCGTGAC TGGG |
| *cfa*-Long GGG mutant gBlock | gBlock gene fragment for *cfa*'-'*lacZ*-Long GGG | TCGCAACTCTCTACTGTTTCTCCATGAGTTGTTAAAG CTACGATAAATATTATGTTTTTACGGGGACAGGATCG TTCCCGACTCACTATGGATAGTCATTTCGGCAAGGGT TCCTCCTTTGGGTCTGTTCTACGTCGGATTATAGACT |

| | | |
|---|---|---|
| | fusion construction | CGCGGTTTTTTCTGCGAGATTTCTCACAAAGCCCAAAAAGCGTCTACGCTGTTTTAAGGTTCTGATCACCGACCAGTGATGGAGAAACTATGAGTTCATCGTGTATAGAAGAAGTCAGTGTACCGGATGACAACTGGTACCGTATCGCCAACGAATTACTTAGCCGTGCCGGTATAGCCATTAACGGTTCTGCCCCGGTCGTTTTACAACGTCGTGACTGG |
| *cfa*-Long GGGGGG mutant gBlock | gBlock gene fragment for *cfa*'-'*lacZ*-Long GGGGGG fusion construction | TCGCAACTCTCTACTGTTTCTCCATGAGTTGTTAAAGCTACGATAAATATTATGTTTTTACGGGGACAGGATCGTTCCCGACTCACTATGGATAGTCATTTCGGCAAGGGTTCCTGGTTTGGGTGTGTTCTACGTCGGATTATAGACTCGCGGTTTTTTCTGCGAGATTTCTCACAAAGCCCAAAAAGCGTCTACGCTGTTTTAAGGTTCTGATCACCGACCAGTGATGGAGAAACTATGAGTTCATCGTGTATAGAAGAAGTCAGTGTACCGGATGACAACTGGTACCGTATCGCCAACGAATTACTTAGCCGTGCCGGTATAGCCATTAACGGTTCTGCCCCGGTCGTTTTACAACGTCGTGACTGGG |
| *Stm cfa*-Long Forward | Forward primer for *S. enterica cfa*'-'*lacZ*-Long fusion construction | ACCTGACGCTTTTTATCGCAACTCTCTACTGTTTCTCCATCGGTTGTTTATATTACGATA |
| *Stm cfa*-Short Forward | Forward primer for *S. enterica cfa*'-'*lacZ*-Short fusion construction | ACCTGACGCTTTTTATCGCAATCTTCTACTCTTTCTCCATAAGGTTCTGATCACCGTCCA |
| *Stm cfa* Reverse | Reverse primer for *S. enterica cfa*'-'*lacZ*-Long and *cfa*'-'*lacZ*- | TAACGCCAGGGTTTTCCCAGTCACGACGTTGTAAAACGACCGGGGCGGAACCATTAATTG |

| | | |
|---|---|---|
| | Short fusion construction | |
| *cfa*-Long transcriptional fusion gBlock | gBlock gene fragment for *cfa*'-*lacZ*-Long transcriptional fusion construction | ACCTGACGCTTTTTATCGCAACTCTCTACTGTTTCTCCATGAGTTGTTAAAGCTACGATAAATATTATGTTTTTACGGGGACAGGATCGTTCCCGACTCACTATGGATAGTCATTTCGGCAAGGGTTCCTCCTTTCCCTCTGTTCTACGTCGGATTATAGACTCGCGGTTTTTTCTGCGAGATTTCTCACAAAGCCCAAAAAGCGTCTACGCTGTTTTAAGGTTCTGATCACCGACCAGTGATGGAGAAACTATGAGTTCATCGTGTATAGAAGAAGTCAGTGTACCGGATGACAACTGGTACCGTATCGCCAACGAATTACTTAGCCGTGCCGGTATAGCCATTAACGGTTCTGCCCCGATTTCACACAGGAAACAGCTATGACCATGATTACGGATTCACTGGCCGTCGTTTTACAACGTCGTGACTGGGAAAACCCTGGCGTTA |
| *cfa*-Short transcriptional fusion gBlock | gBlock gene fragment for *cfa*'-*lacZ*-Short transcriptional fusion construction | ACCTGACGCTTTTTATCGCAACTCTCTACTGTTTCTCCATAAGGTTCTGATCACCGACCAGTGATGGAGAAACTATGAGTTCATCGTGTATAGAAGAAGTCAGTGTACCGGATGACAACTGGTACCGTATCGCCAACGAATTACTTAGCCGTGCCGGTATAGCCATTAACGGTTCTGCCCCGATTTCACACAGGAAACAGCTATGACCATGATTACGGATTCACTGGCCGTCGTTTTACAACGTCGTGACTGGGAAAACCCTGGCGTTA |
| *cfa*-Long +58 transcriptional fusion gBlock | gBlock gene fragment for *cfa*'-*lacZ*-Long +58 transcriptional fusion construction | ACCTGACGCTTTTTATCGCAACTCTCTACTGTTTCTCCATGAGTTGTTAAAGCTACGATAAATATTATGTTTTTACGGGGACAGGATCGTTCCCGACTATTTCACACAGGAAACAGCTATGACCATGATTACGGATTCACTGGCCGTCGTTTTACAACGTCGTGACTGGGAAAACCCTGGCGTTA |
| *cfa*-Long +78 transcriptional fusion gBlock | gBlock gene fragment for *cfa*'-*lacZ*-Long +78 transcriptional | ACCTGACGCTTTTTATCGCAACTCTCTACTGTTTCTCCATGAGTTGTTAAAGCTACGATAAATATTATGTTTTTACGGGGACAGGATCGTTCCCGACTCACTATGGATAGTCATTTCGATTTCACACAGGAAACAGCTATGA |

| | | |
|---|---|---|
| | fusion construction | CCATGATTACGGATTCACTGGCCGTCGTTTTACAAC GTCGTGACTGGGAAAACCCTGGCGTTA |
| *cfa*-Long +98 transcriptional fusion gBlock | gBlock gene fragment for *cfa*'-*lacZ*-Long +98 transcriptional fusion construction | ACCTGACGCTTTTTATCGCAACTCTCTACTGTTTC TCCATGAGTTGTTAAAGCTACGATAAATATTATGT TTTTACGGGGACAGGATCGTTCCCGACTCACTAT GGATAGTCATTTCGGCAAGGGTTCCTCCTTTCCC ATTTCACACAGGAAACAGCTATGACCATGATTAC GGATTCACTGGCCGTCGTTTTACAACGTCGTGAC TGGGAAAACCCTGGCGTTA |
| *cfa*-Long +118 transcriptional fusion gBlock | gBlock gene fragment for *cfa*'-*lacZ*-Long +118 transcriptional fusion construction | ACCTGACGCTTTTTATCGCAACTCTCTACTGTT TCTCCATGAGTTGTTAAAGCTACGATAAATATT ATGTTTTTACGGGGACAGGATCGTTCCCGACT CACTATGGATAGTCATTTCGGCAAGGGTTCCT CCTTTCCCTCTGTTCTACGTCGGATTATATTTC ACACAGGAAACAGCTATGACCATGATTACGGA TTCACTGGCCGTCGTTTTACAACGTCGTGACT GGGAAAACCCTGGCGTTA |
| *cfa*-Long +138 transcriptional fusion gBlock | gBlock gene fragment for *cfa*'-*lacZ*-Long +138 transcriptional fusion construction | ACCTGACGCTTTTTATCGCAACTCTCTACTGTT TCTCCATGAGTTGTTAAAGCTACGATAAATATT ATGTTTTTACGGGGACAGGATCGTTCCCGACT CACTATGGATAGTCATTTCGGCAAGGGTTCCT CCTTTCCCTCTGTTCTACGTCGGATTATAGACT CGCGGTTTTTTCTGCATTTCACACAGGAAACAG CTATGACCATGATTACGGATTCACTGGCCGTCG TTTTACAACGTCGTGACTGGGAAAACCCTGGCG TTA |
| *cfa*-Long +158 transcriptional fusion gBlock | gBlock gene fragment for *cfa*'-*lacZ*-Long +158 transcriptional fusion construction | ACCTGACGCTTTTTATCGCAACTCTCTACTGTTTCT CCATGAGTTGTTAAAGCTACGATAAATATTATGTTT TTACGGGGACAGGATCGTTCCCGACTCACTATGG ATAGTCATTTCGGCAAGGGTTCCTCCTTTCCCTCTG TTCTACGTCGGATTATAGACTCGCGGTTTTTTCTG CGAGATTTCTCACAAAGCCCAATTTCACACAGGAA |

| | | |
|---|---|---|
| | | ACAGCTATGACCATGATTACGGATTCACTGGCCGT CGTTTTACAACGTCGTGACTGGGAAAACCCTGGCG TTA |
| Fusion check Forward | Forward primer for validation of translational fusions | TCGCAACTCTCTACTGTTTCTCCAT |
| Fusion check Reverse | Reverse primer for validation of translational fusions | CCCAGTCACGACGTTGTAAAACGAC |
| *rho* fusion Forward | Forward primer for *rho*'-'*lacZ* fusion construction | ACCTGACGCTTTTTATCGCAACTCTCTACTGTTTCT CCATGACTTCGTATTAAACATACC |
| *rho* fusion Reverse | Reverse primer for *rho*'-'*lacZ* fusion construction | TAACGCCAGCCTTTTCCCAGTCACGACGTTCTAAAAC GACCGGCGTATTCTTTAATTCGG |
| *rho*-R66S check Forward | Forward primer to check for *rho*-R66S mutation | GCTCGTATGCGTAAGCAGGA |
| *rho*-R66S check Reverse | Reverse primer to check for *rho*-R66S mutation | GCGCAAAATAGCGTTCACCT |
| *rne131* check Forward | Forward primer to check for *rne131* mutation | CGCGCCTGTTGTAGCTCCAG |
| *rne131* check Reverse | Reverse primer to check for *rne131* mutation | TTCGCGATTATCGCTGCCTT |

| | | |
|---|---|---|
| *rne3071-cat* linkage Forward | Forward primer to link *cat* gene to *rne3071* allele | TGGGATCGCTGGGGCGGGCATTTTTTTGCCTATTTT GCATGTGTAGGCTGGAGCTGCTTC |
| *rne3071-cat* linkage Reverse | Reverse primer to link *cat* gene to *rne3071* allele | CATATTAAAATTCATCGAATGGCATCCTTGCTAACCA ACACATATGAATATCCTCCTTAG |
| *rne3071* check Forward | Forward primer to check for *rne3071* mutation | AACGCAACTCAGCAGGAAGA |
| *rne3071* check Reverse | Reverse primer to check for *rne3071* mutation | AGACGAACTTCACGCTGGAA |
| *cpxQ* deletion check Forward | Forward primer to check *cpxQ* deletion | TCACGTTCCCAGTAGTAAAC |
| *cpxQ* deletion check Reverse | Reverse primer to check *cpxQ* deletion | ATGCATTAAGCAGCAGGCAA |
| *rydC* deletion check Forward | Forward primer to check *rydC* deletion | CGCGTAAACGTTCCTGAAGG |
| *rydC* deletion check Reverse | Reverse primer to check *rydC* deletion | GGACGTATGGGCAAGGATTA |
| *cfa* 5’UTR Forward | Forward primer for *cfa* 5’UTR RT-qPCR | GTTTTTACGGGGACAGGATCG |
| *cfa* 5’UTR Probe | Probe for *cfa* 5’UTR RT-qPCR | 56-FAM-TCCCGACTC-ZEN-ACTATGGATAGT-3IABkFQ |

| | | |
|---|---|---|
| *cfa* 5'UTR Reverse | Reverse primer for *cfa* 5'UTR RT-qPCR | GGAGGAACCCTTGCCGAAAT |
| *cfa* 5'ORF Forward | Forward primer for *cfa* 5'ORF RT-qPCR | AAGAAGGCTCTTTGGGGTTAGG |
| *cfa* 5'ORF Probe | Probe for *cfa* 5'ORF RT-qPCR | 56-FAM-GATGGCTGG-ZEN-TGGGAATGTGA-3IABkFQ |
| *cfa* 5'ORF Reverse | Reverse primer for *cfa* 5'ORF RT-qPCR | AATGATGGGGGAGTTGGTTCTC |
| *cfa* 3'ORF Forward | Forward primer for *cfa* 3'ORF RT-qPCR | ACGATACCTATTTTGCGGTGGT |
| *cfa* 3'ORF Probe | Probe for *cfa* 3'ORF RT-qPCR | 56-FAM-GGAAGGCAT-ZEN-ATTCCTGCTCCA-3IABkFQ |
| *cfa* 3'ORF Reverse | Reverse primer for *cfa* 3'ORF RT-qPCR | CAGGCAACCGTTCGGAAAAATA |
| *rho* 5'UTR Forward | Forward primer for *rho* 5'UTR RT-qPCR | GCTCGTCACTCAATCCGTCT |
| *rho* 5'UTR Probe | Probe for *rho* 5'UTR RT-qPCR | 56-FAM-TTCTGCGTA-ZEN-CTCTCCTGTGA-3IABkFQ |
| *rho* 5'UTR Reverse | Reverse primer for *rho* 5'UTR RT-qPCR | CATGTCTTTTCGCTGCCTGG |
| *rho* 5'ORF Forward | Forward primer for *rho* 5'ORF RT-qPCR | AACCTCCGCACTGGTGATAC |
| *rho* 5'ORF Probe | Probe for *rho* 5'ORF RT-qPCR | 56-FAM-TCTCTGGTA-ZEN-AGATTCGCCCG-3IABkFQ |

| | | |
|---|---|---|
| *rho* 5'ORF Reverse | Reverse primer for *rho* 5'ORF RT-qPCR | CGTTAACTTTCAGCAGCGCA |
| *rho* 3'ORF Forward | Forward primer for *rho* 3'ORF RT-qPCR | GCAACATGGAACTGCACCTC |
| *gapA* control Forward | Forward primer for *gapA* control RT-qPCR | AAGTGGTTATGACTGGTCCGTC |
| *gapA* control Probe | Probe for *gapA* control RT-qPCR | 56-FAM-ATATGCTGG-ZEN-CCAGGACATCG-3IABkFQ |
| *gapA* control Reverse | Reverse primer for *gapA* control RT-qPCR | CGTTGATAACTTTAGCCAGCGG |
| *cfa* 5'UTR NB probe | Northern blot probe that hybridizes to the *cfa* mRNA 5'UTR | AAATGACTATCCATAGTGAGTCGGG |
| *cfa* CDS NB probe | Northern blot probe that hybridizes to the *cfa* mRNA CDS | GACGCCCACCACGCTTACGTCATAA |
| CpxQ NB probe | Northern blot probe that hybridizes to the CpxQ sRNA | GGGATGGTGTCTATGGCAAGGAAAA |
| RydC NB probe | Northern blot probe that hybridizes to the RydC sRNA | CGAAGAATACGGGTCTACATCGGAA |
| 5S rRNA NB probe | Control northern blot probe that | GTTTCACTTCTGAGTTCGGCATGGGGTCAGGTGGG |

| | | |
|---|---|---|
| | hybridizes to the 5S rRNA | |
| SsrA NB probe | Control northern blot probe that hybridizes to the SsrA tmRNA | ATCCCGTCGAATCCAGAATCAGCCC |
| *cfa*-Long WS mutant gBlock | gBlock gene fragment for *cfa*'-'*lacZ*-Long WS fusion construction | CTGACGCTTTTTATCGCAACTCTCTACTGTTTCTCCATGAGTTGTTAAAGCTACGATAAATATTATGTTTTTACGGGGACAGGATCGTTCCCGACTCACTATGGATAGTCATTTCCCCAAGGGTTCCTCCTTTCCCTCTGTTCTACGTCGGATTATAGACTCGCGGTTTTTTCTGCGAGATTTCTCACAAAGCCCAAAAAGCGTCTACGCTGTTTTAAGGTTCTGATCACCGACCAGTGATGGAGAAACTATGAGTTCATCGTGTATAGAAGAAGTCAGTGTACCGGATGACAACTGGTACCGTATCGCCAACGAATTACTTAGCCGTGCCGGTATAGCCATTAACGGTTCTGCCCCGGTCGTTTTACAACGTCGTGACTGGGAAAACCCTGGCGTTA |
| *cfa*-Long SS mutant gBlock | gBlock gene fragment for *cfa*'-'*lacZ*-Long WS fusion construction | CTGACGCTTTTTATCGCAACTCTCTACTGTTTCTCCATGAGTTGTTAAAGCTACGATAAATATTATGTTTTTACGGGGACAGGATCGTTCCCGACTCACTATGGATAGTCATTTCGGAAAGGGTTCCTCCTTTCCCTCTGTTCTACGTCGGATTATAGACTCGCGGTTTTTTCTGCGAGATTTCTCACAAAGCCCAAAAAGCGTCTACGCTGTTTTAAGGTTCTGATCACCGACCAGTGATGGAGAAACTATGAGTTCATCGTGTATAGAAGAAGTCAGTGTACCGGATGACAACTGGTACCGTATCGCCAACGAATTACTTAGCCGTGCCGGTATAGCCATTAACGGTTCTGCCCCGGTCGTTTTACAACGTCGTGACTGGGAAAACCCTGGCGTTA |
| CpxQ_C10G C11G Forward | Oligo for site-directed mutagenesis of pCpxQ for WS | GATCCAGTATCTTGTTATCC |

| | | |
|---|---|---|
| | compensatory mutation | |
| CpxQ_C10G C11G Reverse | Oligo for site-directed mutagenesis of pCpxQ for WS compensatory mutation | CTTTTCCTTGggATAGACACCATC |
| CpxQ_G9U Forward | Oligo for site-directed mutagenesis of pCpxQ for SS compensatory mutation | ATCCAGTATCTTGTTATCC |
| CpxQ_G9U Reverse | Oligo for site-directed mutagenesis of pCpxQ for SS compensatory mutation | CCTTTTCCTTtCCATAGACAC |
